# The Role of Electron Spin, Chirality, and Charge Dynamics in Promoting the Persistence of Nascent Nucleic Acid-Peptide Complexes

**DOI:** 10.1101/2023.11.16.567399

**Authors:** Pratik Vyas, Kakali Santra, Naupada Preeyanka, Anu Gupta, Orit Weil-Ktorza, Qirong Zhu, Norman Metanis, Jonas Fransson, Liam M. Longo, Ron Naaman

**Affiliations:** Department of Chemical and Biological Physics, Weizmann Institute of Science, Rehovot, 76100, Israel; Institute of Chemistry, The Hebrew University of Jerusalem, Jerusalem 9190401, Israel; Department of Physics and Astronomy, Uppsala University, Uppsala 752 36, Sweden; Earth-Life Science Institute, Institute of Science Tokyo, Tokyo 152-8550, Japan; Blue Marble Space Institute of Science, Seattle, Washington 98104, USA

## Abstract

Primitive nucleic acids and peptides likely collaborated during the earliest stages of biochemistry. What forces drove their interactions, and how did these forces shape the properties of primitive complexes? We investigated the association of two model primordial polypeptides with DNA. Coupling the peptides to a ferromagnetic substrate results in a dependence of the association rate and the extent of DNA binding on the orientation of magnetic moment of the substrate. The DNA binding could be nearly abolished by inverting the orientation of the magnetic field, despite the two polymers having complementary net charges. Inverting the chirality of either the entire peptide or just the connecting cysteine residue inverted the effect of the magnetic moment orientation. These results are attributed to the chiral-induced spin selectivity (CISS) effect, in which molecular chirality and electron spin interact to alter the electric polarizability of the protein. The observation of CISS effects governing simple protein-DNA complexes, suggests that this phenomenon was plausibly operative and potentially significant for primitive biomolecules. A key consequence of the CISS effect is to increase the kinetic stability of primitive protein-nucleic acid complexes. Taken together, our results show how emergent phenomena due to chirality and spin enhance bio-association.

## Introduction

A long-standing debate in evolutionary biology revolves around peptide-first or nucleic acid-first scenarios, questioning whether proteins/peptides (i.e., metabolism) or nucleic acids (i.e., information storage) emerged first in the origins of life. These two elements, however, are not necessarily mutually exclusive – it is likely that they existed in some rudimentary form well before the emergence of the Last Universal Common Ancestor (LUCA).^1–3^ In support of this emerging view is the apparent ease with which peptides and nucleic acids can interact.^3–8^ Model primordial peptides have been shown to readily bind to and forming coacervates with nucleic acids.^9–12^ In some instances, these primordial peptides can even demonstrate elaborate functional interactions, with nucleic acids, that extend from mere binding regimes, such as strand-separation and exchange.^10^ Unlike contemporary binding sites, however, the earliest peptide-nucleic acid complexes were bound less specifically and with weaker bonds. Here we address the question, w*hat forces drove these interactions, that are primitive and yet, evolutionarily relevant, and how did they contribute to the emergence of functional molecular complexes?*

Of particular note is the molecular *polarizability*, the capacity for charge reorganization within a biomolecule, which partially dictates both the kinetics and thermodynamics of binding. The significance of polarizability in protein structure and activity is now increasingly recognized as demonstrated by several experimental and theoretical works.^13–17^ In contemporary proteins, charge reorganization can even extend to sites remote from the active site, a process dubbed charge-reorganization allostery.^18^ These long-range reorganizations are essential for ligand binding and complex formation.^18,19^ It has been demonstrated that the flow of electrons within a biopolymer, and thus its polarizability, is subject to a fundamental phenomenon referred to as chirality-induced spin selectivity (CISS) effect. The CISS effect refers to the experimental observation that the motion of electrons through or within chiral molecules (and materials) depends on their spin state. Which spin is preferred depends on the handedness of the molecule and on the direction of motion of the electrons.^20^ Thus the CISS effect is a direct consequence of molecular chirality and the intrinsic angular momentum, or spin, of the electron. As of now numerous studies have demonstrated CISS-effect in physical, chemical and evolved biological systems.^21–23^ However, whether CISS exists and, if it does, its significance for weakly-interacting primitive peptide-nucleic acid complexes is so far unclear.

Whether CISS was plausibly operative and significant to primitive biomolecular complexes has profound implications for their kinetic stability, as well as for the prebiotic selection of chiral molecules and the potential roles of ferromagnetic minerals. As CISS modulates the rates of electron flow within a biomolecule upon binding, the presence or absence of strong CISS effects suggests distinct regimes of polarizability^24^, and thus differences in the (dis)association process. Moreover, given that CISS is a direct consequence of the chiral centers within biomolecules, it stands to reason that these differences could promote or suppress the recruitment of chiral or achiral molecular forms. Finally, strong CISS effects indicate that, under certain conditions, ferromagnetic minerals such as magnetite could readily alter the binding properties of an electronically coupled peptide or nucleic acid. Thus, while the CISS phenomenon is quantum mechanical in nature, its ramifications extend to the origins of life and biochemistry.^25,26^

To explore the role of CISS in primitive protein-nucleic acid complexes, we leveraged two model primordial peptides: a β-(P-loop)-α motif derived from the ancient P-loop NTPase enzyme family and a simplified helix-hairpin-helix (HhH) motif constructed from just 10 amino acid types.^10,11^ Both peptides interact with nucleic acids and are less than fifty residues long. Notably, the simplified helix-hairpin-helix (HhH) motif is functionally ambidextrous, with both natural and mirror-image peptides readily binding to and forming coacervates with natural nucleic acids.^27^ Upon coupling to a ferromagnet through a reactive cysteine residue, we find that the DNA binding properties of both peptides become highly sensitive to the orientation of magnetic dipole direction of the substrate. Namely, to the direction of electrons’ spin that can be injected into the substrate or from the substrate. Despite charge complementarity between the peptide (positive charge) and the DNA (negative charge), binding can be almost completely abolished upon application of a magnetic moment in the non-preferred orientation. Inverting the chirality of either the entire peptide or just the bridging cysteine residue likewise inverts the effect of the magnetic field. These observations are most reasonably interpreted as evidence for CISS. Finally, using a simple physical model, we show how CISS can augment the kinetic stability of primitive peptide-nucleic complexes to support primordial biopolymer collaboration and accretion. The extended reach of the CISS effect, as demonstrated here, suggests an ancient and enduring preference for molecular worlds rich in chiral centers. Finally, given the ubiquitous nature of peptide motifs used in this study, CISS effects are likely to be wide-spread in modern-protein world.

## Materials and Methods

### DNA and cloning

Synthetic gene fragments encoding P-loop prototypes were sourced from *Twist Biosciences* and cloned into a pET29(+)b expression vector using the standard restriction free cloning method. Similarly, mutant prototypes were generated using standard site-directed mutagenesis via restriction-free cloning with primers obtained from *Integrated DNA Technologies* (IDT).

### Expression and purification of P-loop prototypes

All P-loop prototypes have a C-terminal Trp residue for concentration determination (the prototypes are otherwise devoid of aromatic residues) followed by a 6xHis tag for purification. DNA and amino acid sequences are provided in **Tables S1** and **S2**. Following purification, the yield and purity of purified proteins was assessed by SDS-PAGE. Typically, four peak elution fractions (∼7.5 mL) were pooled together and subjected to two rounds of dialysis (2 h at room temperature followed by overnight dialysis at 4 °C) against buffer containing 50 mM Tris pH 8.0 and 100 mM NaCl. P-loop prototypes generally precipitate during the dialysis step and require an osmolyte such as L-arginine to be resolubilized. The samples were centrifuged and pellets containing the P-loop prototypes were resolubilized in buffer containing 50 mM Tris pH 8.0, 100 mM NaCl, and 1 M L-arginine (“solubilization buffer”). Aliquots of 100-200 µM P-loop prototypes in solubilization buffer were stored at 4 °C and remained soluble and active for at least 10-14 days.

### Total peptide synthesis

The N-αβα prototype with either a C-terminal *L*-Cys (N-αβα_LcysEnd) or *D*-Cys (N-αβα_DcysEnd) residue were purchased from *Synpeptide*. Peptides were dissolved in 10% Acetonitrile + 90% Millipore water according to manufacturer’s instructions to produce 2.5 mM stock solutions. HhH peptides were synthesized in house and their foldedness was assessed (**Supplementary Methods**, **Figures S1** and **S2**). Peptides were divided into aliquots and stored at -20 °C until use.

### Preparation of the ferromagnetic substrate

Ferromagnetic substrates were prepared as described previously. ^28^ An 8 nm Ti adhesive layer was deposited onto a p-type, boron-doped silicon wafer, ⟨100⟩ ± 0.9°. Next, a 100 nm nickel layer, which serves as the ferromagnetic layer, was deposited onto the adhesive-coated Si wafer using an electron beam evaporator. Finally, the Ni layer was coated with a 5 nm layer of gold. The evaporator chamber was kept under high vacuum (<10^-7^ Torr) and ambient temperature during deposition of the metal layers. Substrates were cut into 23 mm × 23 mm squares using a diamond cutter. The substrate pieces were cleaned by boiling in acetone for 10 min, boiling in ethanol for 10 min, and sonicating in water for 1 min to dislodge any broken debris from the surfaces.

### Drop casting

P-loop prototypes were diluted to 100 µM and dialyzed into 50 mM Tris pH 8.0. A ten-fold molar excess of TCEP was added to reduce the C-terminal cysteines. Samples were incubated at room temperature (for biologically purified proteins) or on ice (for synthetic peptides) for 3 hr to ensure complete reduction of all cysteine residues. To remove TCEP, samples were passed through a BioRad *Micro Bio-Spin* P-6 column equilibrated in 50 mM Tris pH 8.0. The protein concentration was measured by spectrophotometry and the final pepetide concentration was adjusted to 80 µM. The HhH peptides were dissolved in 50 mM Tris buffer pH 7.5 and 150 mM NaCl. The concentration of peptide was then adjusted to 20 µM and verified using spectrophotometry. A five-fold molar excess of TCEP was added to reduce disulfide bonds. TCEP was removed as above.

Cleaned substrate (described above) were dried using an N_2_ gas gun one at time and immediately drop cast by placing 30 µL of protein sample in the center of the substrate. The substrates with drop-casted proteins were placed in a Petri dish, sealed with a parafilm, and stored overnight (∼16 hr) at room temperature (biologically produced proteins) or 4 °C (synthetic peptides).

Verification of monolayer formation from overnight incubation of proteins with Au-coated substrates was performed using polarization modulation-infrared reflection-adsorption spectroscopy (PM-IRRAS) and atomic force microscopy (**Supplementary Methods**, **Figures S3**, **S4** and **S5**). FM-substrates were washed three times with 1 mL of 50 mM Tris buffer pH 8.0. A small scratch at the center of each substrate was made using a diamond cutter. The scratch served as a reference point for microscopy experiments.

### dsDNA hybridization

The dsDNA solutions were prepared by mixing the two complementary strands of ssDNA in 0.4 M phosphate buffer pH 7.2. Strands were denatured at 60 °C for 10 min and annealed by cooling to 15 °C at the rate of 1 °C/min. The annealed DNA was passed through a Micro Bio-Spin P-30 column equilibrated in 50 mM Tris pH 8.0 and concentration was measured by spectrophotometry. Finally, DNA samples were adjusted to a concentration of 1 µM duplex DNA.

### Microscopy measurements

Binding kinetics of ssDNA to P-loop prototypes or dsDNA to HhH peptides anchored to ferromagnetic (FM) substrates were monitored using a ZEISS Axio-Observer.Z1 inverted fluorescence widefield microscope. Prepared substrates were then placed on a permanent 0.42 T magnet with the North Pole facing the substrate and secured using adhesive tape. In parallel, 80 µL fluorescently labeled (6-FAM) ssDNA or dsDNA was placed at the center of a 35 mm glass bottom Petri dish (*MAKTEK)*, which was then transferred to the microscope stage. The magnet-substrate assembly was then inverted onto the Petri dish containing the fluorescent DNA solution.

Images were acquired using a 40× air objective lens (LD Plan-Neofluar 40x /0.6). Using a GFP filter, the emitted fluorescence was collected and separated from the excitation beam by two dichroic beam splitters along the optical path. The light was then routed to an avalanche photodiode (APD) for fluorescence imaging. Before initiating test experiments, a control experiment was carried out to focus the z-position at the surface-solution interface. Focusing was done by locating the scratch on the FM-substrate (described above). The z-position was then fixed for all the experiments. Experiments were initiated and fluorescence intensity images were acquired using the *Slidebook* software. Images were collected every 30 s for a total of 15 min. We selected the same 400-pixel x 400-pixel region of each image using the scratch as a reference point. The increase in fluorescence relative to initial fluorescence value (F/F_0_) was calculated and plotted versus time. Each experiment was repeated three times and errors are reported as standard deviations. The entire procedure was then repeated with the South magnet pointing towards the substrate. The F/F_0_ values were fitted to *the one phase association* equation (eq. 1) in GraphPad Prism software to estimate the rate constants.

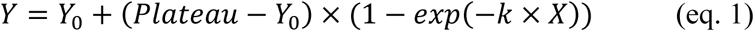

Where *Y*_0_ is the F/F_0_ value at time zero, *Plateau* is the F/F_0_ value at equilibrium, and *k* is the rate constant of the association process expressed in inverse time (s^-1^) units.

### Macroscopic contact potential difference (CPD) measurements

CPD measurements were performed using a commercial Kelvin probe (Delta Phi Besocke, Jülich, Germany) confined inside a Faraday cage under atmospheric pressure. A general Kelvin probe set up includes a metallic probe electrode (gold grid) which is placed near the surface of the sample to form a capacitor. The distance between the probe electrode and the surface of the sample is periodically varied to create a frequency dependent capacitance. An AC voltage is generated across the gap and is proportional to the voltage difference between the sample and the probe electrode. Rather than measuring AC voltage directly, a DC voltage is generally applied to nullify the response, which measures the CPD. The CPD signal is allowed to stabilize before recording.

The substrates were p-doped silicon wafers Si 〈100〉 upon which Ti/Ni/Au 8nm/60nm/8nm with 0.3Å/sec, 0.5Å/sec, and 0.3Å/sec deposition rate at 10^-8^ Torr pressure were grown by an electron beam evaporator. After deposition, the substrates were cleaned with boiling acetone followed by boiling ethanol for 15 minutes each. Immediately after the cleaning, the substrates were dried under nitrogen flow. In parallel, the N-αβα_LcysEnd peptides were incubated with TCEP for reduction of the terminal cysteine as described above. TCEP was removed and the peptides were drop casted on the clean, dry substrates and incubated for 16 hours at 4 °C. Substrates were washed three times with 50 mM Tris pH 8.0, dried with nitrogen gas, and placed on a permanent magnet with either the North polarity facing towards (UP) or away from (DOWN) the substrate. A drop of TC-ssDNA was placed onto the substrates and incubated for 30 min. The dsDNA is then aspirated with a pipette, dried under nitrogen flow, and CPD measurements were performed.

## Results

### Models of primordial protein-DNA complexes

Two families of nucleic-acid binding peptides, both with relevance to early protein evolution, were analyzed (**Table S2**). First, N-αβα and N-βα (**Figures 1A and 1B**) are simplified “prototypes” derived from the core functional fragment of P-Loop NTPase enzymes.^10^ P-Loop prototypes used in this study have been shown to bind cofactors and nucleic acids, promote strand separation and exchange, and weakly catalyze phosphoryl transfer reactions.^10,29^ The profound conservation of the β-(P-loop)-α motif and the broad functional profile of P-Loop prototypes support the hypothesis that these fragments are evolutionary nuclei around which contemporary P-Loop NTPase and other enzyme domains, with the P-loop motif, accreted. ^5,9,30–32^ Second is the *Precursor* peptide (**Figure 1C**), which is derived from an ancestor sequence reconstruction and amino acid deconstruction ^11^ of the ancient and ubiquitous helix-hairpin-helix (HhH) family.^33^ This peptide has been shown to form coacervates with RNA, within which the peptide can dimerize to adopt the (HhH)_2_-Fold.^12^ More recently, it was demonstrated that both the HhH peptide and the (HhH)_2_-Fold are “functionally ambidextrous” -- able to bind RNA and dsDNA of either chirality. ^27^

**Figure 1:**
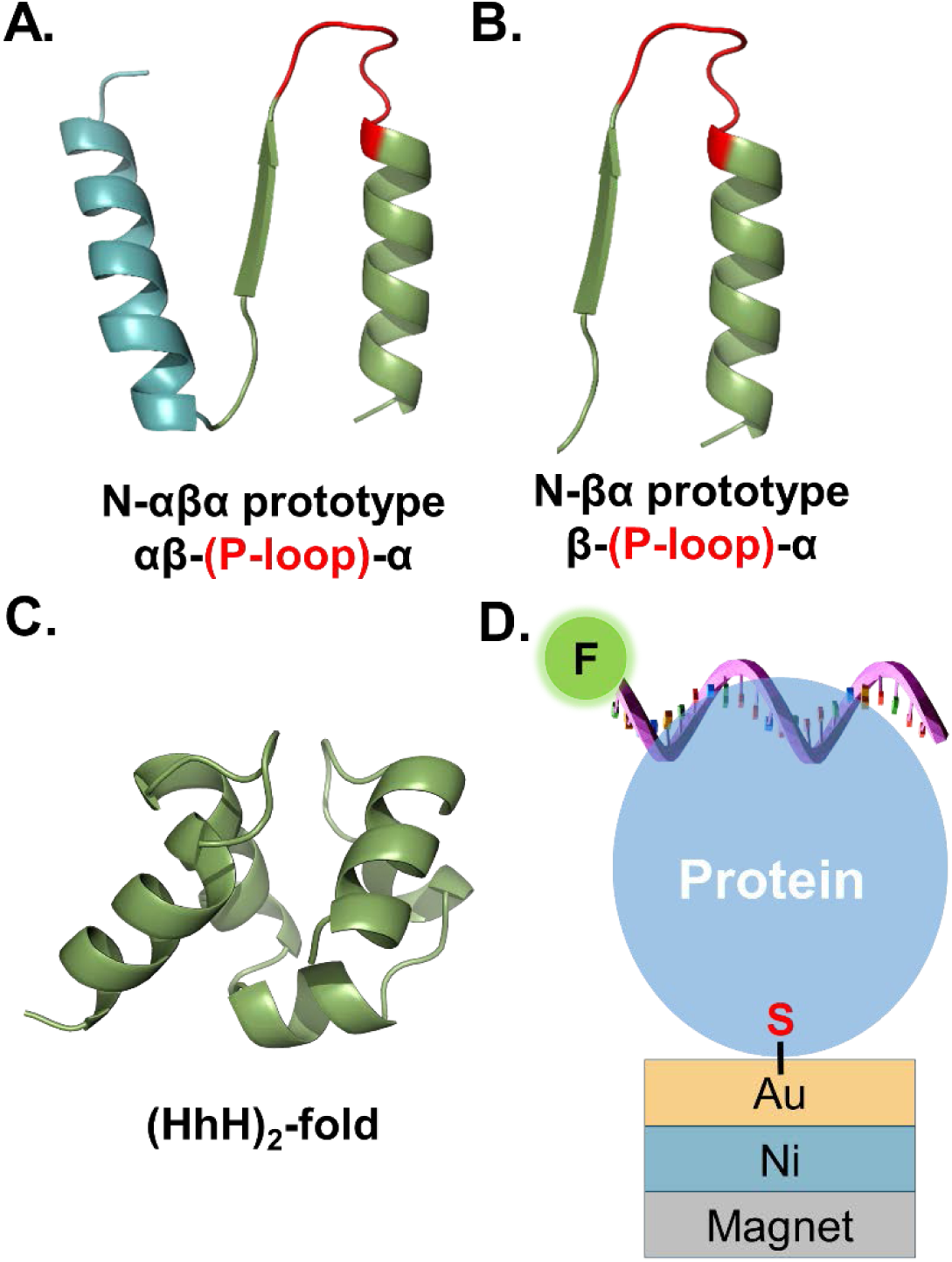
Representative primordial peptide models and experimental setup. **A.** The N-αβα prototype is composed of the ancestral β-(P-loop)- α motif and one additional α-helix at the N-terminus. **B.** The N-βα prototype is composed of only of the ancestral β-(P-loop)-α motif. The structural models in panels A and B were generated using PyMOL and depict the ancestrally inferred β1 and α1 in green and the connecting P-loop in red. The additional α-helix in the N-αβα prototype is shown in cyan. The descriptor below each prototype indicates the the order of secondary structure elements (from N- to C-terminus). The prototypes have a propensity to form dynamic oligomers in solution^9,10,29^; therefore, the monomeric models shown here are only schematic depictions. **C.** AlphaFold2 model of the (HhH)_2_-fold, which is recapitulated upon dimerization of two HhH peptides ^12^ **D.** The experimental setup. Proteins carrying a C-terminal cysteine with a free thiol (-SH) group are adsorbed onto a gold-coated, ferromagnetic nickel layer via an S-Au bond. The ferromagnetic substrates (FM-substrates) with the adsorbed proteins are then placed onto a 0.42T permanent magnet. The entire assembly (magnet + FM-substrates) is transferred to a microscope and binding to fluorescent DNA is monitored. The experiments were carried out with both orientations of magnet (North or South) facing the substrate.

### An experimental setup to detect CISS effects

All molecular binding events induce a dynamic charge redistribution within the participating molecules. Due to the different electrochemical potentials of the molecules that interact, the interaction is equivalent to applying an electric field.^17^ The binding process is mediated in part by the electrical polarizability of the interacting partners, and greater electronic polarizability is generally associated with increased affinity. The electronic polarizability (due to the movement of electrons), in turn, is influenced by the chirality and the spin-state of the migrating electrons i.e. the CISS effect. Which spin is preferred depends on the handedness of the molecule and on the direction of motion of the electrons.^20^ While for one handedness the preferred spin will be always pointing parallel to the electron velocity, for the opposite handedness the preferred spin will always point antiparallel to the electron velocity. A consequence of this effect is that the movement of electrons, in a chiral molecule, generates both a charge-polarization (*i.e.*, a separation of positive and negative charges within the molecule, and spin-polarization (*i.e.*, a preference for electrons with one specific spin, either “up” or “down”) such that one spin has higher density at one electric pole and the other at the opposite pole.^24^ The observed spin polarization typically ranges from 20% to nearly 100% of the moving electrons. ^21^

We measured the CISS effect by coupling one of the binding partners to a reservoir of spin-polarized electrons (**Figure 1D**), as has been done previously.^18,19,21,34^ Briefly, peptides were adsorbed onto a ferromagnetic (FM) substrate that served as a source for spin-polarized charge. To facilitate peptide adsorption, a sole cysteine residue was incorporated at the C-terminus, away from the nucleic-acid binding site. The peptides were then coupled to the surface via the formation of a sulfur-gold bond. Finally, the FM-substrates were attached to one pole of a magnet. The CISS effect predicts that binding affinity and kinetics will strongly depend on the orientation of the magnet because the efficiency of electron flow into or out of the coupled ferromagnet depends on the spin states of the electrons, namely on the direction of magnetization of the substrate. Thus, if the charge reorganization within the protein is spin-dependent, and only electrons with a specific spin are repelled away from the binding site, one magnet orientation will promote nucleic acid binding while the other orientation will reduce it. This effect occurs because the preferred orientation allows electrons to flow into the substrate, increasing the polarizability of the protein and promoting binding. We will return to the question of whether the effects observed in our experiments are due to CISS in the **Discussion**; until then, the observed phenomena will simply be described. Binding was measured via monitoring an increase in fluorescence intensity using a fluorescence microscope. The detailed description of the FM substrate and the binding assay is described in the **Methods.** We will refer to magnet orientations as either UP (North magnet pole facing the FM-substrates) or DOWN (South magnet pole facing the substrates).

### Nucleic acid binding of adsorbed P-loop prototypes is sensitive to magnetic field orientation

The N-αβα prototype binds to fluorescently-labeled ssDNA that is rich in thymine and cytosine nucleotides (TC-ssDNA) with a dissociation constant K_D_ = 0.67 ± 0.16 µM ^10^ (**Figure S6, Tables S3** and **S4**). Adsorption isotherms of TC-ssDNA binding to FM-adsorbed N-αβα prototype exhibit a dependence on the orientation of the magnetic field, with both the rate of binding and the extent of binding increased when the ferromagnet is aligned UP (**Figure 2A**, **Table 1**). Remarkably, the effect of magnetic orientation is so great that binding is nearly abolished when the magnetic field is in the DOWN orientation. However, for ssDNA that is rich in guanines and adenines (GA-ssDNA), which P-loop prototypes bind more weakly with a dissociation constant of K_D_ = 1.70 ± 0.68 µM ^10^ (**Figure S6**; **Table S3** and **S4**), adsorption profiles are similar for both magnetic field orientations (**Figure 2B**).

**Figure 2:**
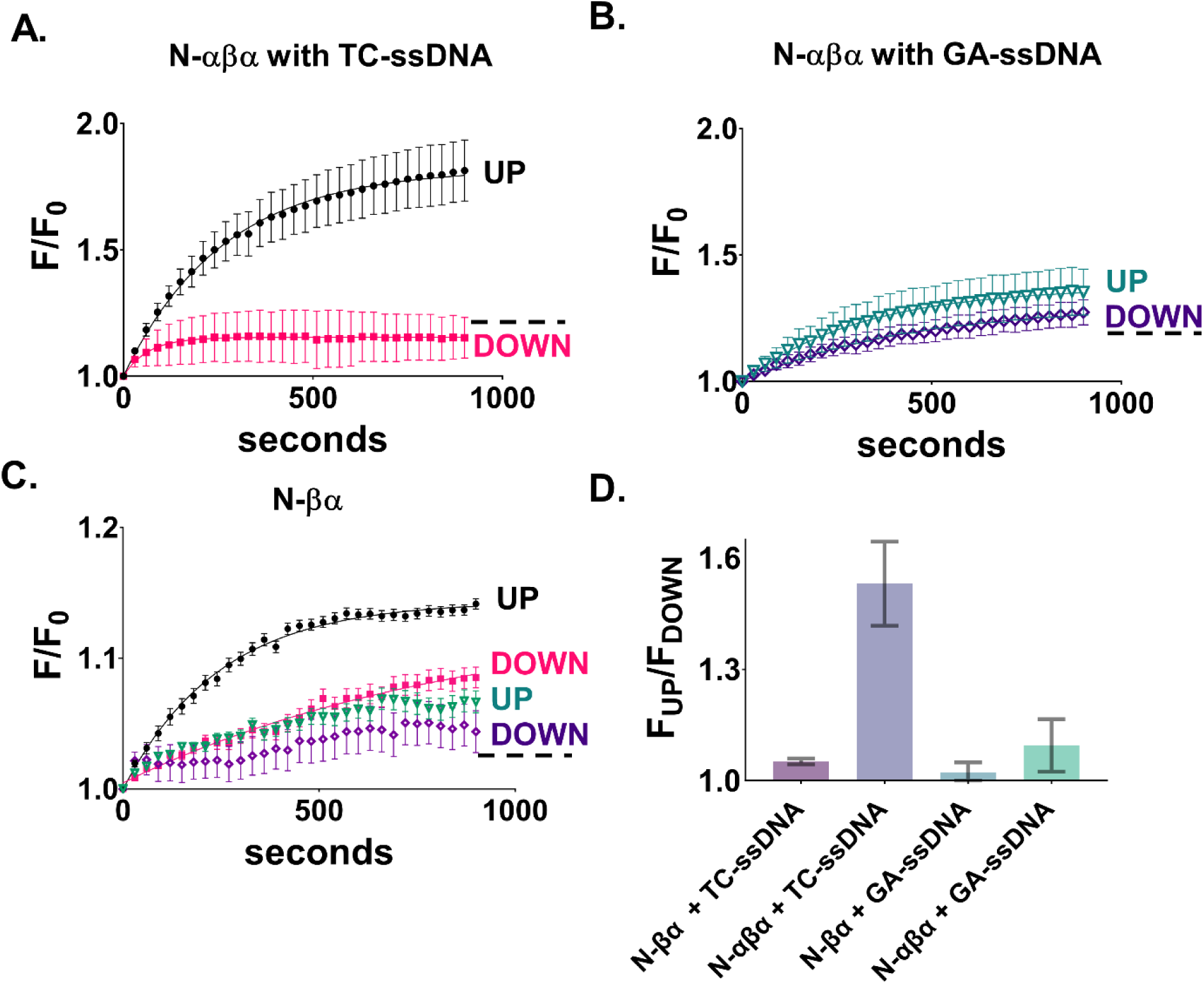
DNA binding to N-αβα and N-βα prototypes is influenced by the orientation of magnet. Adsorption isotherms showing binding of **A.** TC-ssDNA to N-αβα prototype for UP (black trace) and DOWN (pink trace) orientations and **B.** GA-ssDNA to N-αβα prototype for UP (green trace) and DOWN (purple trace) orientations. **C.** Adsorption isotherms showing binding of N-βα prototype to TC-ssDNA for UP (black trace) and DOWN (pink trace) orientations and to GA-ssDNA for UP (green trace) and DOWN (purple trace) orientations. Fits to data in panels **A**, **B**, and **C** were from a standard *one phase association* equation using GraphPad Prism and error bars represent standard deviation from three or four independent measurements. Dotted lines in **A**, **B**, and **C** depict background binding to free fluorescein after nuclease digestion of ssDNA (**Figure S7**) **D.** Relative fluorescence increases. The bar graph displays the relative difference in the endpoint-*F/F_0_* values corresponding to the binding of each prototype (N-αβα or N-βα) to each DNA sequence (TC-ssDNA or GA-ssDNA). Here, the endpoint fluorescence value of the magnet in the UP orientation, was divided by the endpoint fluorescence value of the magnet in the DOWN orientation, for the same DNA-protein pair (F_UP_/F_DOWN_). Error bars represent the standard deviation of three or four independent replicates.

**Table 1:** Apparent rate constants (s^-1^) of DNA binding to adsorbed P-loop prototypes and HhH polypeptides.

|  | TC-ssDNA |  | GA-ssDNA |  |
| --- | --- | --- | --- | --- |
|  | UP | DOWN | UP | DOWN |
| N-αβα | 4 (0.1) X 10<sup>-3</sup><br>*R*<sup>2</sup> = 0.99 | -- | 3 (0.8) X 10<sup>-3</sup><br>*R*<sup>2</sup> = 0.98 | -- |
| N-βα | 4 (0.2) X 10<sup>-3</sup><br>*R*<sup>2</sup> = 0.99 | 1 (0.2) X 10<sup>-3</sup><br>*R*<sup>2</sup> = 0.98 | 3 (0.1) X 10<sup>-3</sup><br>*R*<sup>2</sup> = 0.95 | 0.7 (0.4) X 10<sup>-3</sup><br>*R*<sup>2</sup> = 0.88 |
| N-αβα_LcysEnd | 6 (2) X 10<sup>-3</sup><br>*R*<sup>2</sup> = 0.97 | -- |  |  |
| N-αβα_DcysEnd | -- | 8 (4) X 10<sup>-3</sup><br>*R*<sup>2</sup> = 0.98 |  |  |
|  | 12-bp dsDNA |  |  |  |
|  | UP | DOWN |  |  |
| L-Precursor | 1.9 (0.1) X 10<sup>-2</sup><br>*R*<sup>2</sup> = 0.82 | -- |  |  |
| D-Precursor | -- | 1.8 (0.5) X 10<sup>-2</sup><br>*R*<sup>2</sup> = 0.76 |  |  |

The behavior of the N-βα prototype, which lacks the N-terminal α-helix (**Figure 1B**), is similar to that of N-αβα: binding to TC-ssDNA is favored when the magnet is in the UP orientation and binding to GA-ssDNA is weak and insensitive to magnetic field orientation (**Figure 2C** and **2D**). However, the magnetic field effect is notably smaller for the N-βα prototype than it is for the N-αβα prototype, likely owing to the overall weaker binding of this construct (**Figure 2D**). To verify that the observed increase in fluorescence is not an artifact of the N-αβα prototype interacting with the fluorophore on the DNA, but rather a consequence of prototype-DNA binding, the adsorption of nuclease-digested TC-ssDNA to the N-αβα prototype-coated substrate was measured. As this sample contains only the free fluorophore and nucleotides, a time-dependent increase in fluorescence would reflect binding to the fluorophore. The fluorescence amplitude of the binding isotherms and the effect of the magnetic field, however, are both small consistent with minimal background binding. (**Figure S7**).

### Reciprocal inversion of chirality and magnetic orientation effects

P-loop prototypes have a C-terminal cysteine residue that serves as a linker between the protein and the magnetized FM-substrate, serving as the site for electron injection. N-αβα prototypes that have opposite handedness of the C-terminal cysteine residue, but are otherwise composed of *L*-amino acids, were chemically synthesized. N-αβα_LcysEnd and N-αβα_DcysEnd bear a *L*-Cys and *D*-Cys residue, respectively, at the C-terminus (**Table S2**). While the N-αβα_LcysEnd peptide favored the UP orientation – like the *E. coli*-expressed N-αβα prototype (**Figure 3A**) – the N-αβα_DcysEnd peptide favored the DOWN orientation (**Figure 3B**). As before, the non-preferred orientation of the magnetic field blocks the binding down to the level of the background (**Figure S8**).

**Figure 3:**
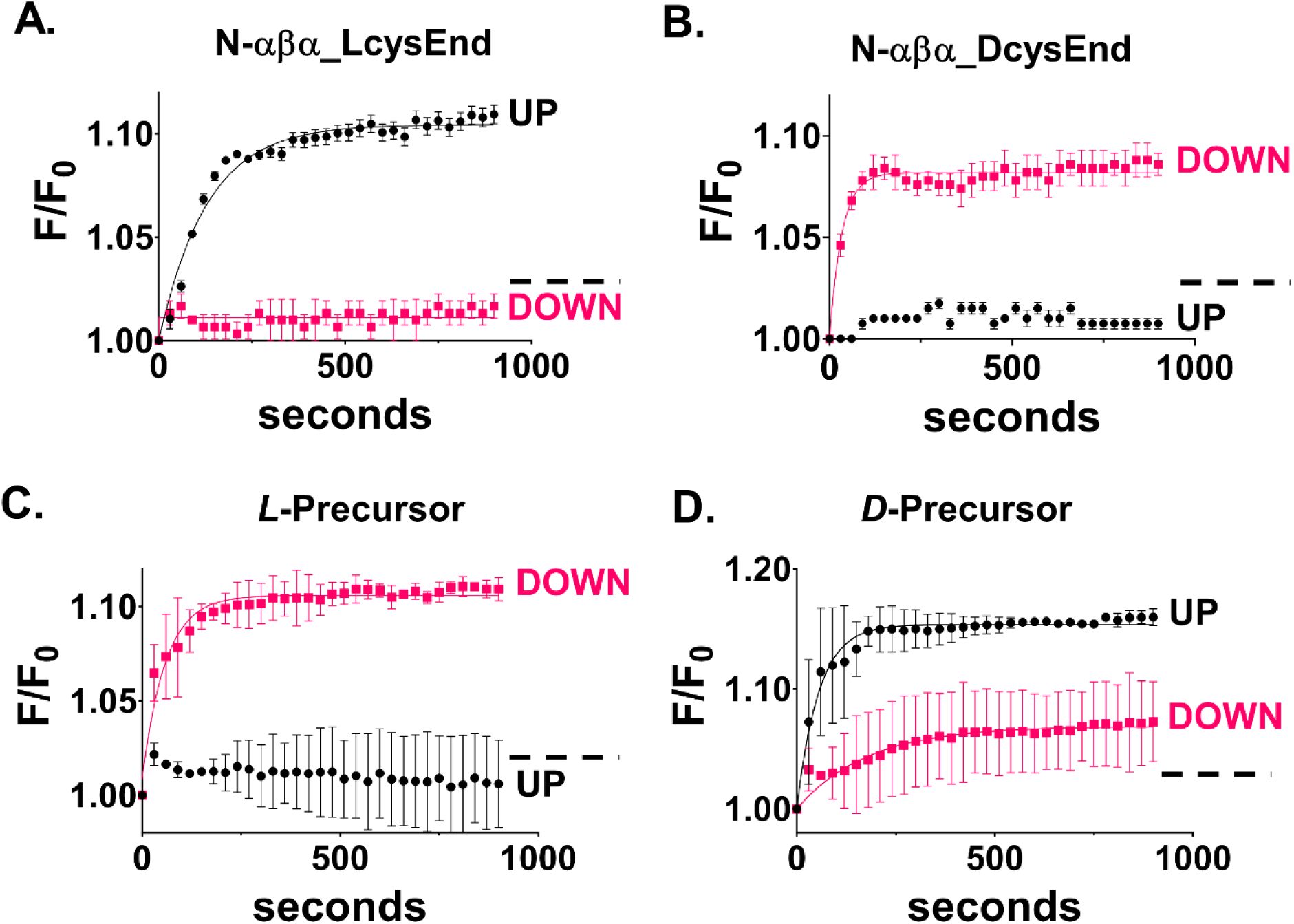
Handedness effect on the magnetism-dependent association. The influence of handedness of synthetic P-loop prototypes and HhH polypeptides on binding to TC-ssDNA or 12-bp dsDNA, respectively, for both magnet orientations. **A.** Adsorption isotherms of TC-ssDNA binding to N-αβα anchored to a ferromagnetic surface via *L*-cysteine (N-αβα_LcysEnd). **B.** Same as A but with N-αβα anchored via *D*-cysteine (N-αβα_DcysEnd). **C.** Adsorption isotherms of 12-bp dsDNA binding to the Precursor-Arg peptide, which is comprised of a single HhH motif, and for which with all amino acids are in the *L*-form (*L*-Precursor) **D.** Same as **C** but with all amino acids in the *D*-form (*D*-Precursor). Error bars represent standard deviation from three or four independent measurements. Dotted lines in **A**, **B**, **C** and **D** depict background binding of free fluorescein.

We note that the amplitude of the fluorescence-increase for reactions with biologically purified proteins (**Figure 2**) is larger than that for chemically synthesized proteins (**Figure 3**). The greater fluorescence signal for biologically produced proteins is reproducible upon multiple batches of purification (**Figure S9**) and may relate to differences in oligomeric state. In addition, the histidine tag used for purification of the biologically produced samples may change the oligomerization state ^29^, resulting in greater binding to DNA but also higher non-specific interactions with the hydrophobic fluorescent dye. Nonetheless, previous studies have shown that the histidine tag does not modulate the function of P-loop prototypes. ^9,10^ Crucially, the relative preference for the UP orientation for both N-αβα prototype samples remain unchanged.

Finally, reciprocal inversion of magnetic field orientation preference and chirality was tested on the HhH peptide binding to dsDNA (**Figures 3C** and **3D**). To this end, we test two HhH peptides that are composed exclusively of either *L*- or *D*-amino acids, ^27^ thus inverting the entire handedness of the protein construct (including their DNA-binding sites) and assuming that the binding mode is largely unchanged. Whereas the *L*-Precursor peptide enjoys increased affinity for dsDNA when the substrate is magnetized DOWN, the *D-*Precursor peptide has an inverted response with increased affinity for dsDNA when the substrate is magnetized UP. Note that while the preferred magnetic orientation is not conserved between the P-Loop NTPase and HhH-derived peptides, both types of peptides exhibit a clear orientation preference that can be inverted by inverting the chirality of either the linker (P-Loop NTPase) or the entire peptide (HhH). In all cases, the binding rates for *L*- and *D*-peptides are similar for the favored magnet orientations (**Table 1**), suggesting that the actual rate of binding under ideal conditions is the same for the two enantiomers and the electron spin is not involved in the binding site itself.

Lastly, as an independent verification of the results obtained using fluorescence microscopy, we performed contact potential difference (CPD) studies with a Kelvin probe ^35^ (**Figure 4**; refer to **Methods** for experimental details). Briefly, Kelvin probe experiments allow measurement of the *work-function,* the threshold energy required to remove an electron from a solid surface. These measurements can confirm the extent of ssDNA binding for different magnetization directions by measuring the electric field between a sample and a metallic probe. The setup includes a metallic probe electrode (gold grid) placed near the surface of the sample to form a capacitor (**Figure 4A**). The distance between the probe electrode and the surface of the sample is periodically varied to create a frequency-dependent capacitance. As a result of the CPD, an AC voltage is generated across the gap that is proportional to the difference in the work-function between the sample and the probe electrode. A DC voltage is applied to nullify the AC voltage. A more negative value indicates that the adsorbate reduced the work-function more significantly.

**Figure 4.**
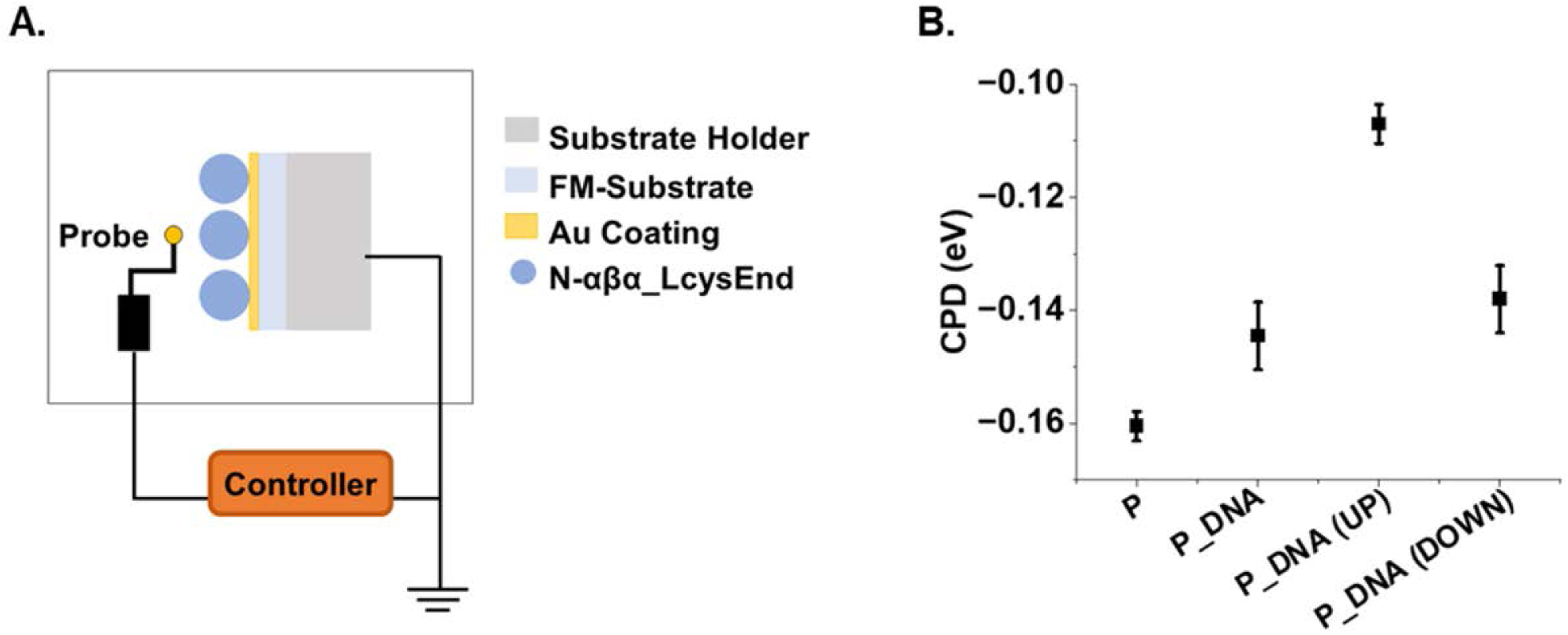
**A.** Schematic of the Kelvin probe setup. **B.** The measured CPD values for chemically synthesized N-αβα. CPD (eV) values were measured for ‘Protein-only’ (P), adsorbed protein without incubation with DNA; for (P_DNA)’, adsorbed protein incubated with DNA with the substrate not magnetized; ‘Protein_DNA (UP)’, adsorbed protein incubated with DNA with magnet facing towards the protein; and Protein_DNA (DOWN), adsorbed protein incubated with DNA with magnet facing away from the protein. Error bars represent the standard deviation from four independent experiments.

As a representative construct of the proteins described in this study, chemically synthesized, N-αβα_LcysEnd prototype was adsorbed onto FM-substrates and binding was measured as described in the **Methods** section. The UP orientation exhibited a less negative work-function (*i.e.*, less negative CPD value and higher work function) than the down orientation (**Figure 4B**), indicating that more ssDNA is adsorbed onto the protein-coated. In comparison, for the surfaces coated with protein only, the CPD values are the most negative, consistent with the least material present on the surface. For surfaces covered with protein and ssDNA bound in the absence of an external magnet field, the CPD values are similar to the DOWN orientation. Taken together, the Kelvin probe experiment corroborates the results from fluorescence microscopy.

## Discussion

### The CISS effect accounts for the sensitivity of nucleic acid binding to magnetic field orientation

The magnetization-dependent dependent binding functions suggests that the peptides are spin-selective. Numerous studies have evidenced that the CISS effect underlies various physical, chemical and biological systems. ^21–23^ Therefore, prima facie, given its fundamental nature, one might assume that the CISS effect contributed to primitive peptide-nucleic acid systems as well. However, that CISS exists in ancient biomolecular interactions is far from obvious. Previous studies demonstrating the CISS effect, involved standard and evolved biological (or physical and chemical) systems that demonstrate clear, robust spin-dependent interactions facilitated by well-defined chiral structures and specific binding affinities. ^18,19,34^ Typically, these systems exhibit strong, highly specific interactions at active sites that enhance the CISS effect. In contrast, primordial proteins are unlikely to have a defined tertiary and quaternary structure as their modern counterparts, possess have weak binding affinities, with promiscuous active sites that may engage in non-specific interactions, thus potentially hindering the CISS-effect.

The experimental set-up show in Figure 1 allows us to determine spin-selectivity of the peptides and the resulting CISS effect. Since the orientation of ferromagnet dictates the spin orientation that is injected into the adsorbed proteins, the results suggest that P-loop prototypes and HhH peptides demonstrate electron spin-selective DNA binding-functions. It is important to note that the role of the ferromagnetic substrate is only to polarize the spin of the electrons within the Ni/Au substrate and its magnetic field has no direct effect on the protein-DNA binding. Applying an external magnetic field, without having spin polarized substrate, does not affect the DNA-protein binding. The extent of the spin-selective effect is highly sensitive to the strength of ligand binding – with only a ∼2.5-fold difference in K_D_ resulting in a near-complete loss of the CISS effect for some protein-ligand pairs. More specifically, in the case of P-loop prototypes binding to the TC-rich sequence or of the HhH peptides to dsDNA, the amount of charge that is reorganized is large, and hence charge has to be spread all over the system. However, for GA-rich DNA where the binding to the P-loop prototype is weaker, only a small amount of charge reorganization occurs and, consequently, the charge reorganization is short range and does not result in a significant charge exchange with the substrate (**Figure 2A** and **2B**).

The strongest evidence in favor of the CISS effect comes from the experiments involving chiral-inversion (**Figure 3**). Specifically, flipping the handedness of a single terminal residue of P-loop prototypes inverts the spin-preference for nucleic-acid binding *i.e.* DOWN orientation is favored for prototypes with terminal *D-*cysteine, whereas UP orientation is favored for prototypes *L-* cysteine terminus (**Figures 3A** and **3B**). Schematically, we suggest that when the linker is *L*-Cys it shows the same preference as the native protein, and thus provides a path for the electrons with the preferred spin to move into the ferromagnetic substrate when magnetized UP. For *D*-Cys, on the other hand, electrons with the same spin cannot pass efficiently through the linker and instead accumulate between the linker and the binding site to the DNA, reducing the effective polarizability of the protein. However, if the magnet is pointing DOWN, electrons from the *D*-Cys can now be displaced into the substrate, hence the electrons repelled from the binding site can now be inserted (after their spin is flipped) into the *D*-Cys. The fact that the relative spin orientations between the substrate and the linker, and not the linker and the remainder of the protein, dominates the handedness effect, suggests a clear separation of timescales that will form the basis of future work along with studies to quantify the spin-polarization effects shown in Figures 2, 3 and 4.

The observed spin selectivity can, in principle, be a result of the spin-dependent charge polarizability within the proteins or due to the spin-dependent interaction between the DNA and the protein. The question of which mechanism is the dominant one was resolved by studying the spin-dependent association with HhH peptides that are composed exclusively *L*- or *D*-amino acids, thus inverting the entire handedness of the protein construct (**Figure 3D** and **3E**). The rates of binding for the *L*- and *D*-HhH peptides are similar for the favored spins (**Table 1**), suggesting that while spins influence the polarizability of the peptides, resulting in inverted spin preferences for the two enantiomers, the actual rate of binding, once polarizability is maximized, is the same for the two enantiomers. Consequently, the interaction between the DNA and the peptide is not spin dependent, and the observed effect is instead the result of spin-dependent charge polarizability within the protein. This finding supports the conclusion that the interaction is predominantly electrostatic without significant selectivity to the chiral structure.

Overall, inverting the chirality of either the terminal cysteine or the entire peptide construct changes the preference for magnet orientation, underscoring the role of the peptides in mediating spin-polarized DNA-binding functions. These results also rule out any spurious surface effects, such as magnetostriction *i.e.* a change in shape of ferromagnetic materials when subjected to magnetic field, for the UP vs DOWN orientation. The CISS effect is consistent across two distinct ancient peptides, various sequences of DNA, types of DNA (single stranded vs double stranded DNA) and are corroborated by an independent Kelvin-probe method (**Figure 4**).

### Evolutionary implications of magnetization-dependent activity

Mineral-organic interfaces may have played a crucial role in the functionalization of life and bio-catalysis in the early stages of the earth. ^36–38^ Binding to minerals has also been shown to preserve protein sequences and accordingly biominerals have been proposed to be excellent source of ancient protein sequences. ^39^ Minerals and rock surfaces were also likely to have served as compartments and scaffolds for prebiotic reactions and protocells. ^40,41^ Nickel deposits (the ferromagnetic layer used in our studies) abundant on the Archean earth, ^42^ played a key role in biological processes, including enzymatic reactions.^43,44^ Iron is the most abundant element in the earth’s crust and it is postulated to be ubiquitous in the primordial world. ^45–48^ Beyond acting as protective scaffolds, these magnetic minerals may have created unique spin-polarized environments. In a plausible scenario, primordial peptides may have functioned when adsorbed onto mineral interfaces, as previously described, or onto magnetite, which can provide source of spin-polarized electrons upon UV irradiation,^25,26^ thereby conferring chiral preference to the adsorbed peptides and, as a consequence, facilitating the transition from heterochiral to homochiral biomolecules. Foremost, given that DNA-binding functions of primordial polypeptides can be promoted or abrogated depending on the substrate magnetization, the functioning of peptides adsorbed to magnetic minerals provides a means to augment binding interactions and on-rates between ancient biomolecular interactors.

It is true that the magnetic field used in this study to observe the spin-polarization is much stronger than the magnetic fields present on early earth. However, it has been shown that similar spin-polarizations can be achieved by superparamagnetic magnetite particles, even when they are formed under weak magnetic fields similar to that of the early earth, due to their remnant magnetizations ^26,49^. Furthermore, since the effect we describe results from potential spin-exchange with the substrate, it is the degree of spin-alignment at the protein-substrate (or mineral) interface that is crucial, rather than rather than earth’s magnetic field. In other words, the magnetization of naturally occurring minerals is the primary driving force for achieving spin alignment at the protein-substrate (or mineral) interface, rather than the strength of Earth’s magnetic field. ^26^

### Significance of electron-spin polarizability for contemporary protein/enzyme systems?

As described below, the importance of spin selectivity is rationalized by a schematic (**Figure 5**). When the DNA approaches the protein adsorbed on the ferromagnet, it induces polarization of the charge in the protein. Here we refer not to the charge localized on specific functional groups, but rather to charge that can be delocalized. This polarization is related to the high frequency polarizability of the protein, or in other terms, to the protein’s high frequency dielectric constant. Since the DNA is negatively charged, when it approaches the protein electrons are repelled from the binding site (**Figures 5A** and **B**). Because of the chirality of the protein, the charge rearrangement depends on the electrons’ spin.^24^ Namely, electrons with one spin orientation move more efficiently through the protein. If the substrate is magnetized so that the electrons with the preferred spin can be injected into it, then the electrons withdrawn from the binding site can move into the ferromagnet, thereby the effective polarizability of the protein increases. In the specific case of *L*-proteins, this means that since the preferred spin for the electrons’ motion is the spin antiparallel to the electrons’ velocity, the ferromagnet should be aligned with its North pole pointing towards the protein (namely its majority spin is aligned away from the protein) and therefore the electrons from the protein can be injected into the minority states of the ferromagnet (**Figure 5A**).

**Figure 5:**
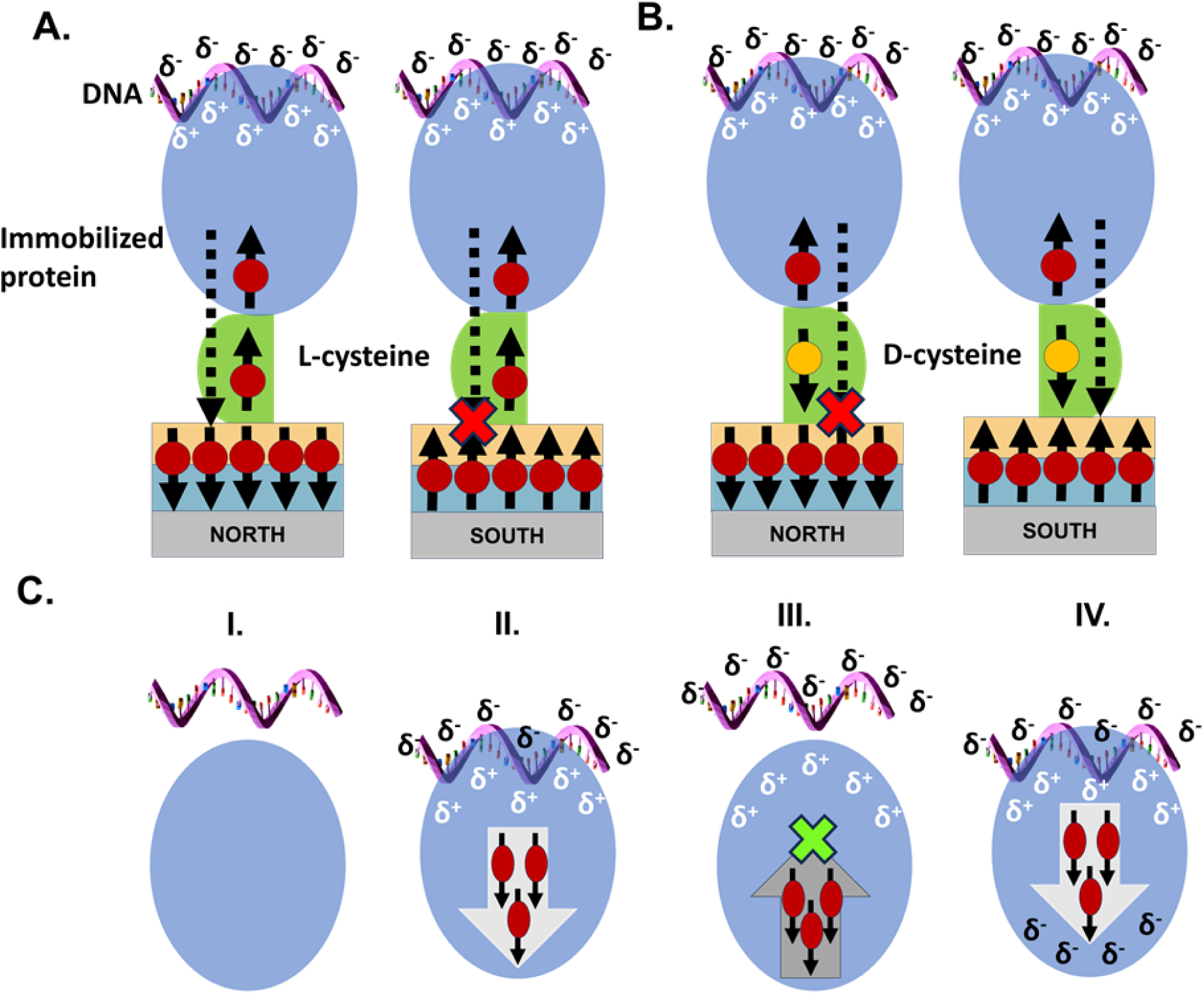
**A.** Schematic representation of the mechanism of the effect of spin on the protein-DNA interaction. When the spin on the ferromagnet is pointing opposite to the momentary spin of the charge at the interface of the adsorbed proteins (representative spins and electrons migrating trough the protein are shown as black arrow and red ball respectively), charge flows more efficiently between the protein and FM-substrate (shown as dotted arrow penetrating the substrate). This scenario manifests when the North magnet is facing the FM-substrate. When the South magnet is facing the FM-substrate, the spins on the protein and the substrate are pointing the same direction resulting in inefficient electron-flow. **B.** Effect of the *D*-cysteine linker on protein-DNA interaction. The terminal *D*-cysteine linker, shown in green, which has the opposing chirality to the rest of the protein, flips the direction of spins transmitted through the protein (black arrow, yellow ball). Consequently, the opposite magnet polarity is required to achieve electron-flow, relative to the schematic in A. **C.** Model describing the role of electrons spin-related polarizability in the docking of DNA on the protein. Reaction scheme describes the dynamic interaction of a protein molecule with a DNA molecule to form transient (Protein-DNA)’ complex (charge-reorganization step) and the final energetically favored (Protein-DNA) configuration (binding step). **I**. The system before DNA-protein interaction. **II**. The approaching DNA repels electrons with specific spins away from the binding zone. **III**. Upon retraction of the DNA from the protein, the electrons that were withdrawn are not able to flow back because they have the “wrong” spin direction. **IV**. The final docking that results in binding of the DNA to the protein.

The phosphate binding motifs used in this study are ubiquitous in present-day protein world. ^5,9,11,29,30^ The results therefore suggest that CISS is at play in contemporary protein interactions. This spin dependency of the association is surprising, and the question is if it serves any function in biological systems. In what follows we will present a possible explanation for the importance of the spin. It is well accepted that the “docking” of substrates/ligands on enzymes/proteins is not an abrupt single-step event but a rather a dynamic process, influenced by the conformational dynamics of the participating biomolecules, involving multiple ‘binding-unbinding’ events, until the minimum-energy configuration is achieved. ^50–54^ Upon the first encounter of the DNA and the protein, electrostatic interactions induce charge-reorganization in the protein, *i.e.* due to the approaching negatively charged DNA, electrons are repelled from the binding site in the protein, giving rise to positive charge polarization (**Figure 5C, panel II**). As a result of the CISS effect, electrons with spin parallel to their velocity move faster than those with spin aligned antiparallel to their velocity. When the DNA molecules transiently “unbind” from the protein, some of the charge will move back into the binding site, however for these electrons to move back, their spins must flip. This spin-flipping is a relatively slow process that has a lifetime of hundreds of nanoseconds, leading to a delay in the charge returning to its original state (**Figure 5C, panel III**). As a result, the internal charge dynamics of the protein is quenched and the polarization, therefore, remains positive. Thus, the DNA molecules are attracted back towards the protein. This dynamic enhances their binding affinity to the protein and prevents the complete unbinding of DNA before a minimal energy conformation is reached (**Figure 5C, panel IV**). It is important to appreciate that interaction of protein with any binding-partner will involve charge-reorganization within the protein, due to the different electrochemical potentials of the protein and the binding-partner. Hence, the mechanism described above should be relevant to all association processes, in which the objects are chiral.

Given the current state of art methods, it is not possible to simulate a fully realistic, theoretical model to support the schematic shown in Figure 5. Yet, we have attempted simplified quantum simulations, as described in previous works, ^55,56^ to model the dynamics of the charge density using model chiral and achiral systems. The simulations show significant spin polarization in chiral molecules, while almost negligible effect for achiral molecules. Further, for chiral molecules electrical and spin polarization are related, whereas these are independent properties for achiral molecules (**Supplementary Text:** *Quantum mechanical simulations*; **Figure S10**).

## Conclusion

We have demonstrated that the association between model primordial polypeptides and DNA depends on electron spin. The spin dependence is proven by the dependence of the association on the direction of the spin polarization in the ferromagnetic substrate. We also showed that the effect of the spin depends on the handedness of the protein. For *L* protein the enhancement is with one spin while for *D* protein it is with the opposite spins. We performed model calculations aiming at demonstrating the proposed mechanism for the spin effect. The experimental results, supported by the schematic illustration and a simplified quantum calculation, clearly demonstrates the difference between chiral and achiral systems and highlights the role of the electron’s spin in making this difference. The spin effect arises due to spin-selective polarizability of the protein due to the chiral-induced spin-selectivity (CISS) effect.

The CISS effect links homochirality (be it *L* or *D*) to the spin-dependent polarization of electrons which, in turn, influences internal charge dynamics that affect their function. We propose that the spin selectivity enhances the binding rate between protein and substrate by elongating the relaxation time of the ‘chiral-induced electric dipole’ moment in the protein. If true, spin selectivity may have promoted the emergence of homochirality by augmenting and modulating binding kinetics in weakly-interacting, ancient protein-nucleic acid systems. Given the ubiquity of the motifs studied, our study suggests that CISS effect is wide-spread in contemporary biomolecular systems. In conclusion, our study provides evidence of CISS effect regulating functions in tangible, primordial protein systems, thus highlighting an underappreciated consequence of homochirality that may have facilitated cooperation amongst nascent biological systems.

## Acknowledgments

We thank Dr. Reinat Nevo for help with microscopy experiments. RN acknowledges partial support from the US-AFSOR Grant FA9550-21-1-0418. JF acknowledges support from Vetenskapsrådet and Stiftelsen Olle Engkvist Byggmästare. NM acknowledges the support of the Israel Science Foundation (1388/22). PV acknowledges financial support from the Feinberg Graduate School, Weizmann Institute of Science. The DNA icons in Figures 1 and 5 were obtained from *bioicons.com*.

## Data and materials availability

The authors declare that the data supporting the findings of this study are available within the paper and its Supplementary Information files. Should any raw data files be needed in another format they are available from the corresponding authors upon reasonable request. Source data are provided with this paper.

