## Supplementary Information file for "The Role of Electron Spin, Chirality, and Charge Dynamics in Promoting the Persistence of Nascent Nucleic Acid-Peptide Complexes"

**Affiliations**

**This PDF file includes:**

Supplementary Text: Quantum mechanical simulations
Supplementary Text: Materials and Methods
Supplementary Figures: S1 to S10
Supplementary Tables S1 to S4
References (1 to 5)

### Supplementary Text

*Quantum mechanical simulations*

For testing if the proposed mechanism of spin-dependent charge reorganization shown in Figure 5 (main text) can in principle take place, we performed simplified quantum simulations to model the dynamics of the charge density with a model system, comprising of a chain of $\mathbb{M}$ sites distributed in helical (chiral) or a zig-zag (achiral) shape (**Figure S10, panel A**). We shall refer to these model systems as the chiral and achiral molecules. Each site may hold an electron at the energy $\varepsilon_m$ and is coupled to its nearest and next-nearest neighbor by both elastic $(t_0, \lambda_0)$ and inelastic $(t_1, \lambda_1)$ components, where the inelastic components account for nuclear vibrations that are coupled to the electronic charge $(t_1)$ and to the electronic spin through the vibrationally enhanced spin-orbit coupling $(\lambda_1)$. The chiral or achiral structures are controlled by the next-nearest neighbor interaction. The effect of the perturbing charge on the chiral system is provided by an additional external electron, illustrated by the magenta ball in **Figure S10, panel A**, where the green line between this charge and the system signifies the (time-varying) electrostatic coupling between the DNA and the protein. The details of setting up the molecule are described later in **Materials and Methods** (*Simulation Methododogy*) and in Refs. [1,2].

The simulations of the charge dynamics in the chiral and achiral chain are undertaken by varying the coupling between the external charge and the system harmonically as $U_0(1 + \cos v_0 t)$, where $U_0$ defines the maximum coupling strength, $v_0$ the frequency of the variations, and $t$ is the time. The method used for the simulations of the dynamics is described in the Supplementary Information and in Ref. 2.

In Figure S10, panel B, we have plotted the density of electron states for the chiral and achiral molecules. The spectrum, which is confined below the chemical potential, zero vertical line, should be considered as the set of occupied orbitals whereas the unoccupied orbitals are assumed to be located at energies beyond 4 eV. It should be noticed that the densities for the two molecules are identical.

By the time-dependent variations in the local charge conditions near one of the edges of the system that models the molecule (protein), the internal charge of the molecules forced to react in response to these changes. As a measure of the variations, we calculate the electric polarization vector $P = (Px, Py, Pz)$ of the system with respect to its center of mass. In both types of molecules, there is a distinctive time-dependence imposed on the charge polarization (**Figure S10C** and **S10D**), where largest vector component of $P$ is plotted as function of time. Here, we compare the dynamical responses for frequencies between 1 MHz and 10 THz, and the responses are plotted on the same unit less time axis to enable a comparison.

First, one can notice that the overall polarization is stronger in the chiral than in the achiral structure, both in amplitude and mean value. Second, it can be seen that the time-dependent variations tend to be strongly suppressed with in increasing frequency in the chiral structure compared to the achiral one. In the chiral molecule, there is a clear trend of decreasing amplitude in the temporal response the higher the frequency, a trend which cannot be discerned in the achiral molecule.

In **Figure S10E** and **S10F** we also plot the associated time-dependent spin-polarization for the chiral and achiral molecules, respectively, and there are two striking differences between the induced spin-polarizations. First, while there is a significant dynamical spin-polarization in the chiral molecules, there is none at all in the achiral (notice that the vertical scale is in units of $10^{-15}$). Second, for the chiral molecule, also the spin-polarization tends to become increasingly time-independent with increasing frequency, whereas there are hardly any changes in the achiral molecule.

The results from the calculations clearly point to that the electric and spin polarizations are intimately related in chiral molecules, while these quantities may be considered as independent in achiral. This conclusion can be drawn from three observations. First, since the densities of electron states in the two molecules are identical (**Figure S10B**), one can effectively exclude the possibility that the responses to the time-dependent perturbation would be a density effect. Second, since the chiral molecule develops a spin-polarization whereas the achiral does not, in response to the time-dependent perturbation, we can deduce that the induced spin-polarization is the key quantity that is distinct between the two molecules. Third, the fact that the spin-polarization in the chiral molecule becomes increasingly time-independent at a non-vanishing mean value with increasing frequency (**Figure S10E**), indicates that charges with different spins accumulate in different spatial locations and that there is a spatial imbalance built up through this differentiation of the spin. This spatial imbalance can, hence, be detected as an electric polarization (**Figure S10C**).

***Materials and Methods***

**Synthesis and purification of peptides with the HhH motif**

*Synthesis:* As described in the ref. [3], *L*-Precursor and *D*-Precursor peptides, were synthesized on 2-Chlorotrityl-resin (loading 0.3 mmol/g, on a 0.25 mmol scale) on automated peptide synthesizer. Peptides were deprotected and cleaved to give 1356 mg of crude *L*-Precursor and 405 mg of crude *D*-Precursor.

*Purification:* 200 mg of crude *L*-Precursor and 100 mg of crude *D*-Precursor peptides were purified by RP-HPLC (XSelect C18 column, 5 µm, 130 Å, 30 × 250 mm) using a gradient of 30-60% B over 42 min to give pure *L*-Precursor (68 mg, 34% yield) and pure *D*-Precursor (26 mg, 26% yield). The HPLC analysis (**Figure S1**) was carried out on a C4 analytical column.

**Circular Dichroism (CD) characterization of the HhH peptides**

Secondary structural characteristics of *L*- and *D*-Precursor peptides was assessed by CD measurements. The CD spectroscopy measurements were performed by using a using a nitrogen purged Chirascan™-Plus spectrometer, (Applied Photophysics, UK), with a thermoelectrically controlled single cell holder. The measurements were conducted at 1 s per point, 1 nm step size, and 1 nm bandwidth. The optical path of the quartz cuvette used was 0.1 cm. CD spectra of *L*- and *D*-Precursor indicates that the peptides are largely unfolded and are mirror images of each other (**Figure S2**)

**Polarization modulation-infrared reflection-adsorption spectroscopy (PM-IRRAS) characterization of peptides.**

PM-IRRAS was carried out to ensure overnight (16 hr.) incubation of FM-substrates with N-αβα prototype and HhH peptides results in a monolayer formation. For these experiments we used a 100 nm Au-coated silicon substrate. Spectrum was obtained for overnight adsorbed proteins by accumulating 2000 scans with the samples mounted at Brewster angle of incidence of 80 on a Nicolet 6700 FTIR with a PEM-90 photoelastic modulator. The spectra of peptide monolayers showed two characteristic peaks at 1670 and 1540 $cm^{-1}$, typical of amide-I (stretching mode of the CO bond) and amide-II (N−H in- plane bending mode and C−N stretching mode), respectively (**Figure S3** and **S4**).

**Atomic Force Microscopy (AFM) for topography analysis.**

AFM topography of N-αβα prototype and L-*Precursor* peptide monolayer on gold coated Si surface was scanned by Bruker-AFM MultiMode 8 in tapping mode with sharp silicon tips (radius = 7 nm, spring constant = 2 N/m, Model AC240TS-R3 from Oxford instruments) (**Figure S5**).

**Simulation methodology**

The simulations are based on the time-dependent Green function for the molecule in presence of a time-dependent disturbance. The molecule is modeled by the Hamiltonian $\mathcal{H} = \Psi^\dagger(H_0 + H_1)\Psi + \mathcal{H}_{ph}$, where $\Psi = \{\psi_m\}_{m=1}^{\mathbb{M}}$ is a column vector of the $\mathbb{M}$ spinors $\psi_m = (\psi_{m\uparrow}\psi_{m\downarrow})^t$, and $H_0$ and $H_1$ are the non-interacting and interacting contributions to the spectrum, while $\mathcal{H}_{ph} = \sum_\mu \omega_\mu b_\mu^\dagger b_\mu$ represents the collective nuclear vibrations distributed over the modes $\mu$ with energy $\omega_\mu$ which are created and annihilated by the operators $b_\mu^\dagger$ and $b_\mu$, respectively. Here, the first contribution to $\mathcal{H}$ can be written

$$H_0 = \left\{\epsilon_m\delta_{mn} + \sum_{s=\pm1}(-t_0\delta_{nm+s} + i\lambda_0\boldsymbol{v}_m^{(s)}\cdot\boldsymbol{\sigma}\delta_{nm+2s})\right\}_{mn=1}^{\mathbb{M}},$$

where $\epsilon_m$ denotes the ons-site HOMO level, $t_0$ is the nearest-neighbor hopping, and $i\lambda\boldsymbol{v}_m^{(s)}\cdot\boldsymbol{\sigma}$ is the next-nearest neighbor hopping. The latter component connects the chirality $\boldsymbol{v}_m^{(s)}$ with the spin-orbit interaction $i\lambda_0$. The second contribution to $\mathcal{H}$ is given by

$$H_1 = \left\{\sum_{s=\pm1}(-t_1\delta_{nm+s} + i\lambda_1\boldsymbol{v}_m^{(s)}\cdot\boldsymbol{\sigma}\delta_{nm+2s})\sum_\mu(b_\mu + b_\mu^\dagger)\right\}_{mn},$$

where $t_1$ and $\lambda_1$ define corresponding vibration assisted nearest and next-nearest neighbor interactions, through the coupling to the nuclear displacement operator $\sum_\mu(b_\mu + b_\mu^\dagger)$.

The molecule is coupled to a time-independent external charge $\psi_0^\dagger\psi_0$ via the charge-charge interaction $V(t)\psi_0^\dagger\psi_0\psi_1^\dagger\psi_1$, with the time-dependent coupling parameter $V(t) = U_0(1 + cos\hbar v_0 t)$, where $v_0$ defines the frequency of the coupling.

The molecular charge and spin distributions are calculated using non-equilibrium Green functions, $\mathbb{G}(t,t') = \{\boldsymbol{G}_{mn}(t,t')\}_{mn}$, where each matrix element $\boldsymbol{G}_{mn}(t,t') = (-i)\langle T\psi_m(t)\psi_n^\dagger(t')\rangle$ is a

$2 \times 2$-matrix propagator. The site resolved time-dependent charge and spin densities are given by $\langle n_m(t)\rangle = (-i)sp\boldsymbol{G}^{<}_{mm}(t,t)/2$ and $\langle \boldsymbol{s}_m(t)\rangle = (-i)sp\boldsymbol{\sigma}\boldsymbol{G}^{<}_{mm}(t,t)/4$, respectively. The equation of motion for $\mathbb{G}(t,t')$ is approximated by

$$(i\partial_t - H_0)\mathbb{G}(t,t') = \delta(t-t') + \langle\psi_0^\dagger\psi_0\rangle(t)\mathbb{V}(t)\mathbb{G}(t,t') + \int \boldsymbol{\Sigma}(t,\tau)\mathbb{G}(\tau,t')d\tau,$$

where $\mathbb{V}(t)$ is the matrix coupling the external charge $\langle\psi_0^\dagger\psi_0\rangle(t)$ to $\boldsymbol{G}_{11}(t,t^{'})$ and $\boldsymbol{\Sigma}$ is the self-energy caused by the interactions with the nuclear vibrations.

Here, we define the Green function $\mathbb{G}_0$ to include the interactions between electrons and nuclear vibrations, that is,

$$(i\partial_t - H_0)\mathbb{G}_0(t,t') = \delta(t-t') + \int \boldsymbol{\Sigma}(t,\tau)\mathbb{G}(\tau,t')d\tau,$$

which is time-independent. Hence, we may write

$$\mathbb{G}_0(z) = (z - H_0 - \boldsymbol{\Sigma}(z))^{-1}.$$

Using this formulation, the time-dependent Green function is provided by the expression

$$\mathbb{G}(t,t') = \mathbb{G}_0(t,t') + \int \mathbb{G}_0(t,\tau)\mathbb{V}_0(\tau)\mathbb{G}(\tau,t')d\tau,$$

where $\mathbb{V}_0(t) = \langle\psi_0^\dagger\psi_0\rangle(t)\mathbb{V}(t)$. For computational purposes, this expression is approximated by

$$\mathbb{G}(t,t') \approx \mathbb{G}_0(t,t') + \int \mathbb{G}_0(t,\tau)\mathbb{V}_0(\tau)\mathbb{G}_0(\tau,t')d\tau.$$

The lesser Green function $\mathbb{G}^<(t,t^{'})$ is, then, given by the expression

$$\mathbb{G}^<(t,t^{'}) \approx \mathbb{G}_0^<(t,t^{'}) + \int (\mathbb{G}_0^r(t,\tau)\mathbb{V}_0(\tau)\mathbb{G}_0^<(\tau,t^{'}) + \mathbb{G}_0^<(t,\tau)\mathbb{V}_0(\tau)\mathbb{G}_0^a(\tau,t^{'}))d\tau.$$

*Supplementary Figures*

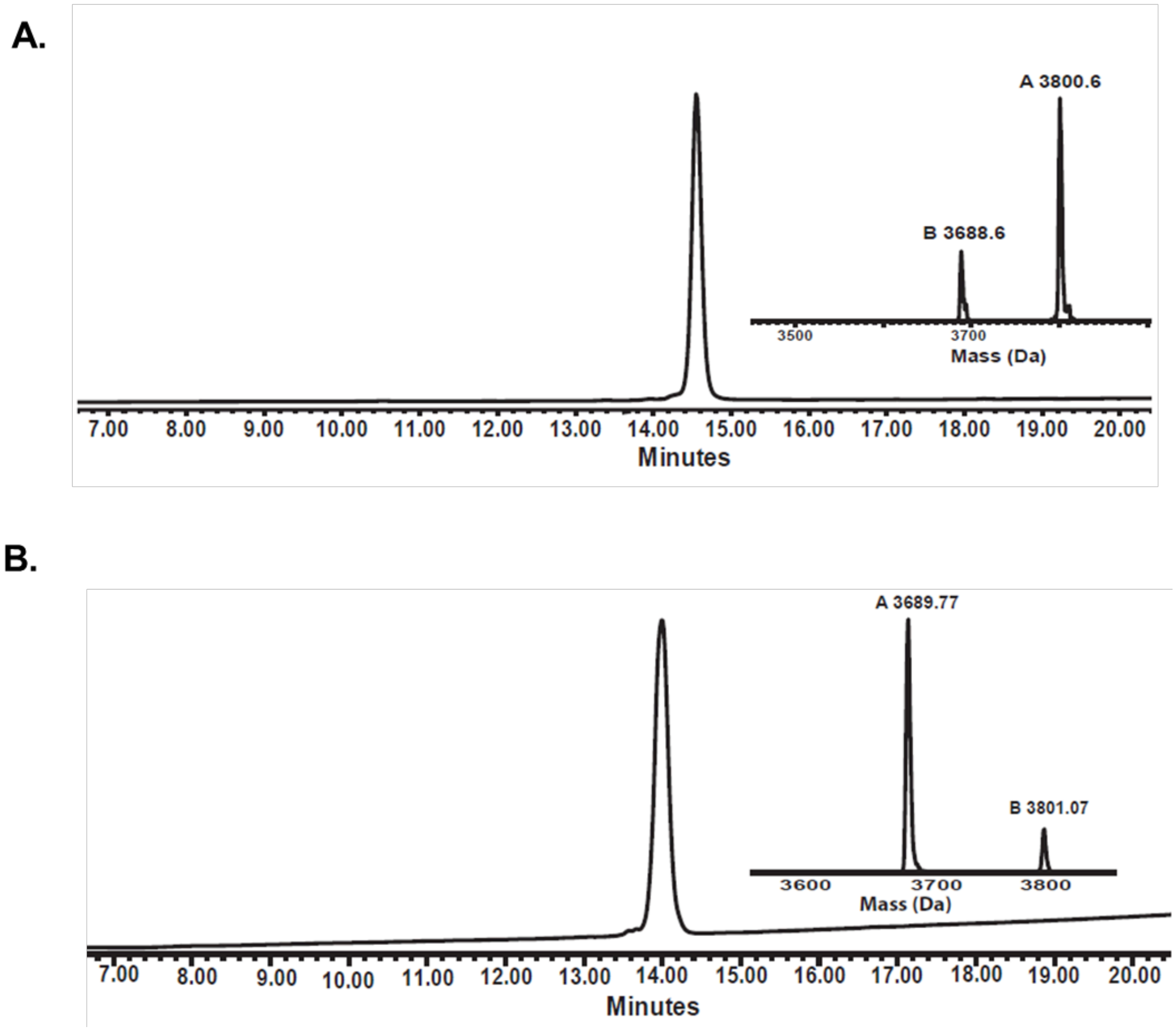


**Figure S1:** HPLC chromatograms and ESI-MS for **A.** purified *L*-Precursor, with the inset showing the corresponding mass (calc. 3689.3 Da; obs. 3688.6 Da, [M+TFA] 3800.6 Da) and **B.** purified *D*-Precursor, with the inset showing the corresponding mass (calc. 3689.35 Da; obs. 3689.77 Da, [M+TFA] 3801.07 Da).

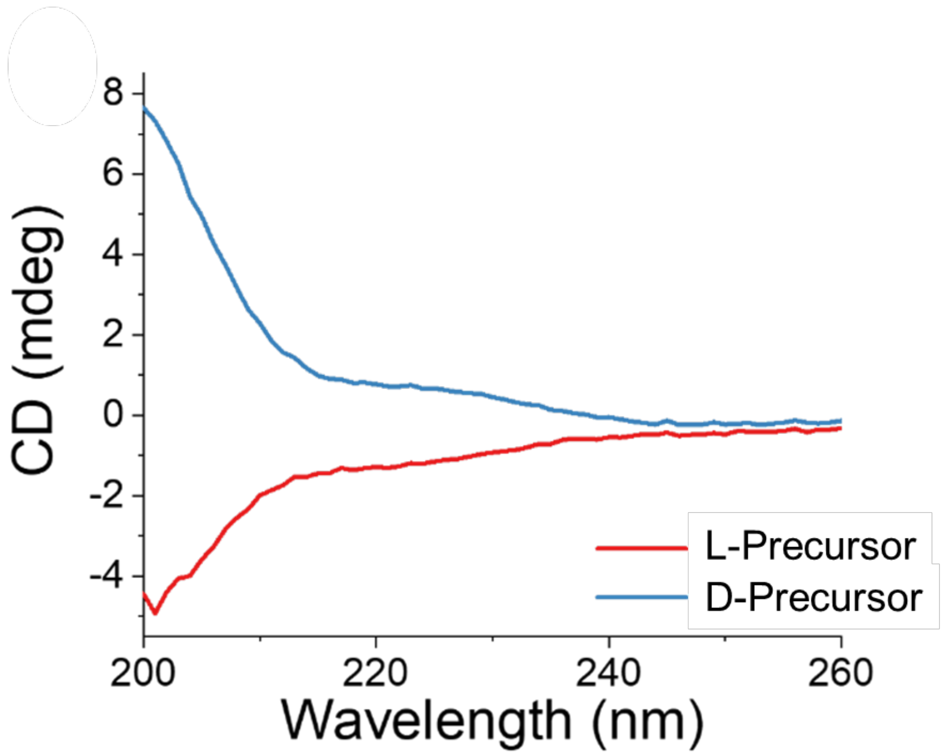


**Figure S2:** CD spectra of *L*- and *D*-Precursor peptides in 50 mM Tris buffer with 150 mM NaCl showing a predominantly unfolded conformation.

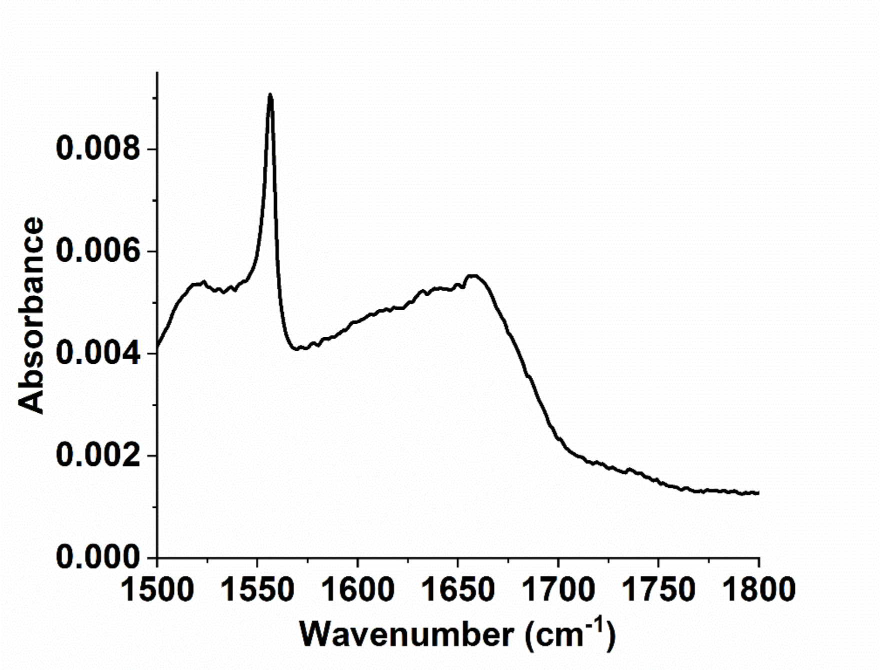


**Figure S3:** A representative PM-IRRAS spectra of the N-αβα prototype incubated for 16 hr. on Au substrate. Here 80 µM peptide was used to grow the monolayer. The peak around 1670 and 1540 $cm^{-1}$ correspond to the characteristic C=O stretching N-H in plane bending mode and C−N stretching mode vibrations, commonly named as amide-I and amide-II peak.

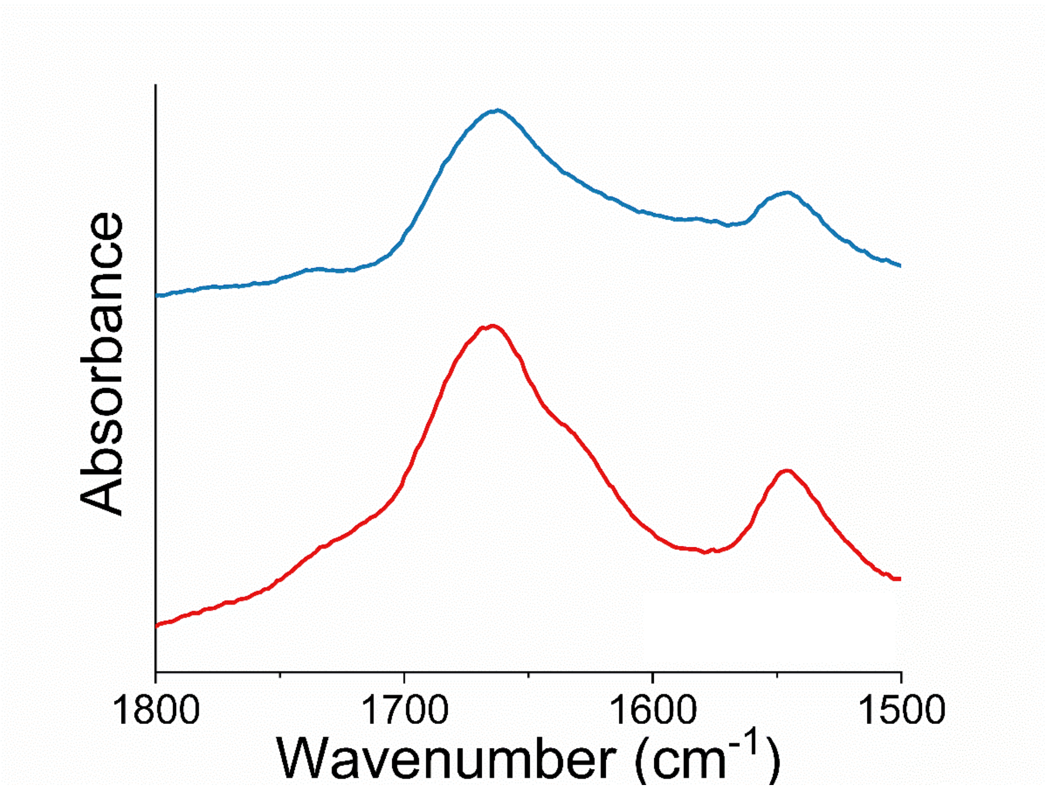


**Figure S4:** PMIRRAS spectra of *L*-Precursor (red) and *D*-Precursor (blue) peptide incubated for 16 h on Au substrate. Here 20 µM peptide was used to grow the monolayer. The peak around 1670 and 1540 $cm^{-1}$ correspond to the characteristic C=O stretching N-H in plane bending mode and C−N stretching mode vibrations, commonly named as amide-I and amide-II peak.

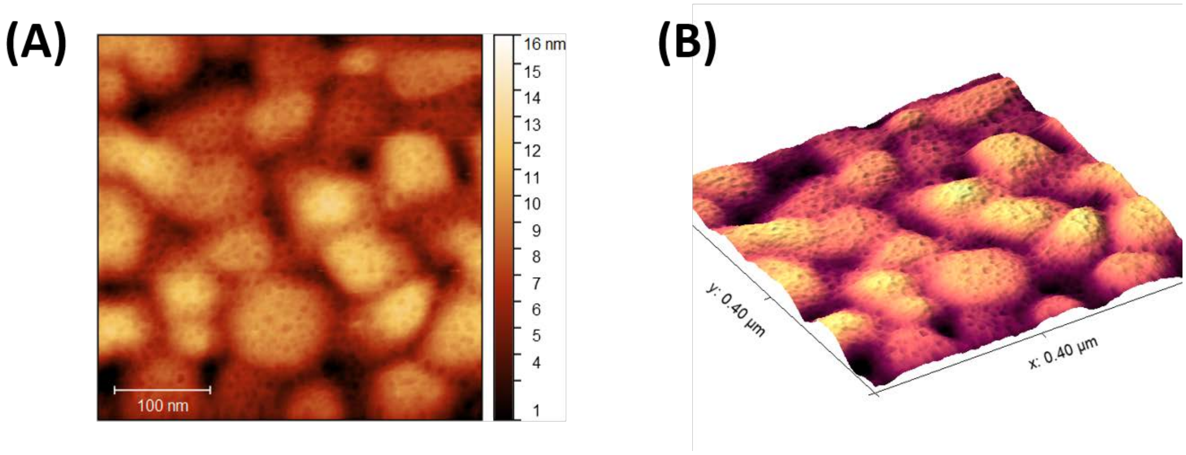


**Figure S5:** A representative AFM topographic **A**. 2D and **B.** 3D image of the *L*-Precursor HhH peptide monolayer on Au surface.

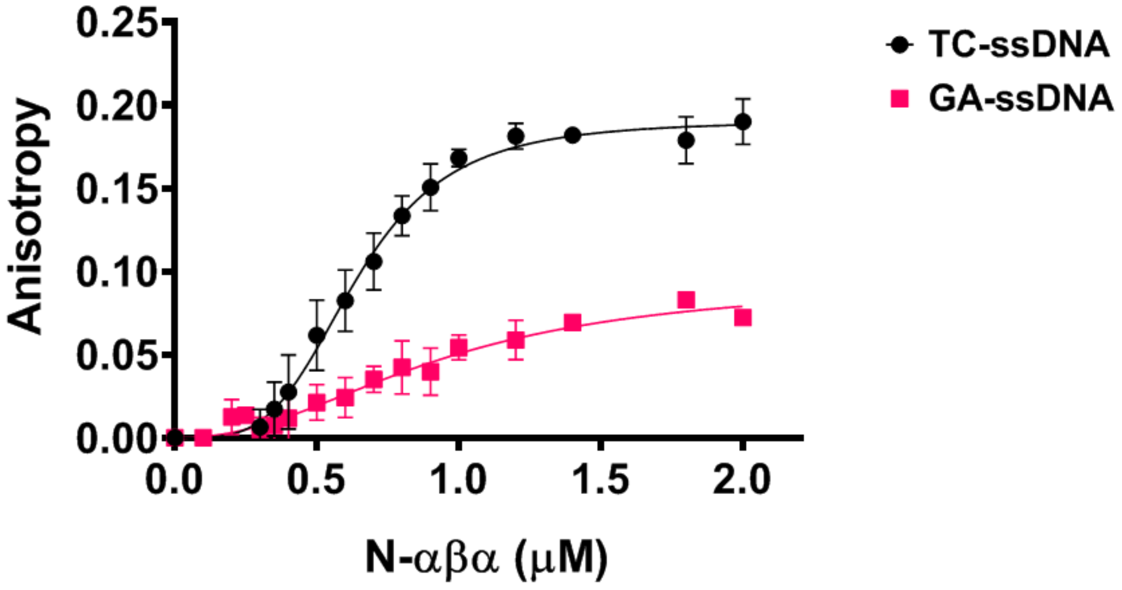


**Figure S6:** Binding of N-αβα prototype to DNA measured by fluorescence anisotropy**.** Shown is the normalized anisotropy signal. Increase in anisotropy indicates binding. Figure is reproduced from ref. [4]. Shown here is fluorescence anisotropy plot of varying concentrations of the N-αβα prototype with TC- and GA-ssDNA oligos (**Supplementary Table S1**). Shown are average values from four to eight independent experiments with vertical error bars representing the SD values. For details on methods see ref. [4]. The KD values are listed in table S4.

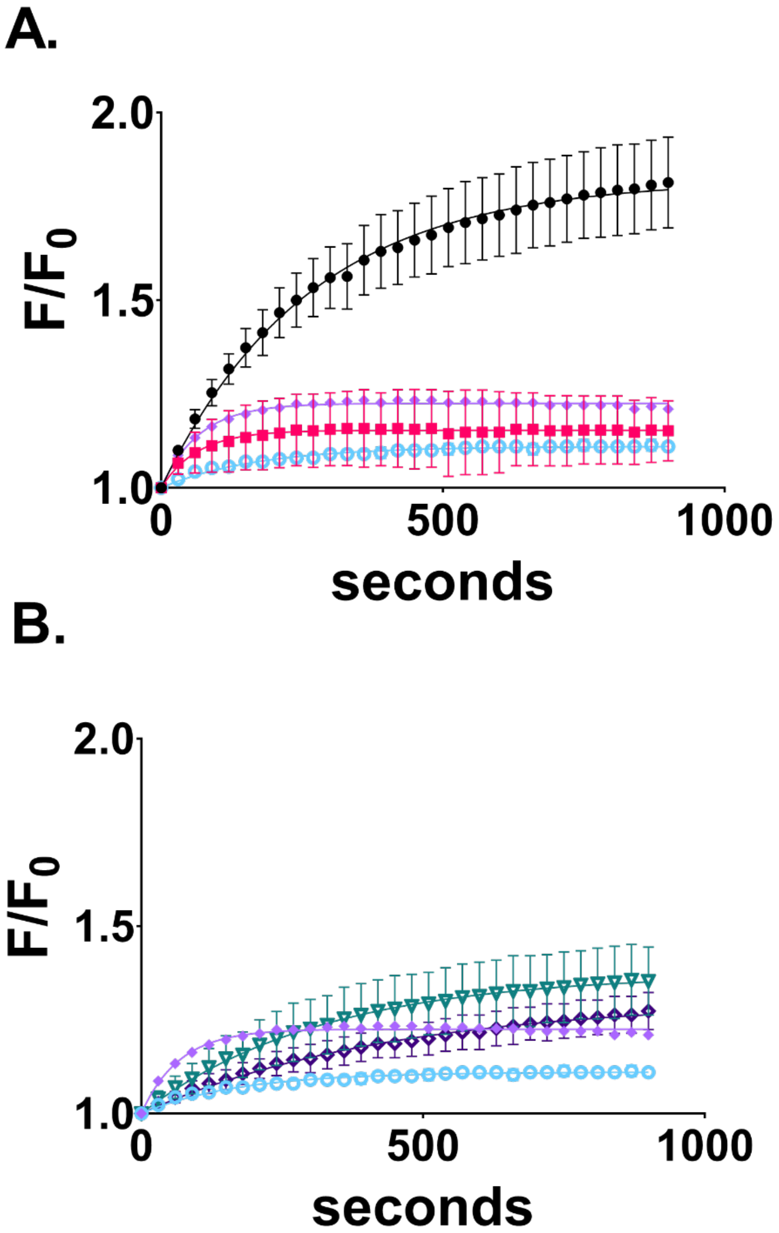


**Figure S7: Comparative adsorption profiles of N-αβα prototype to DNA and fluorescein dye. A.** Adsorption isotherms of N-αβα binding to TC-ssDNA and fluorescein dye for UP and DOWN orientations of the magnet. *TC-ssDNA + UP (black)*, *TC-ssDNA + DOWN (pink)*, *fluorescein + UP (purple)*, *fluorescein + DOWN (blue)* **B.** Adsorption isotherms of N-αβα binding to GA-ssDNA and fluorescein dye for UP and DOWN orientations of the magnet. *GA-ssDNA + UP (green)*, *GA-ssDNA + DOWN (violet)*, *fluorescein + UP (purple)*, *fluorescein + DOWN (blue)* Data were plotted as $F/F_0$ vs seconds and fitted to standard one-phase association equation using GraphPad Prism software. Error bars represent standard deviation from three to four independent measurements. Adsorption curves for TC-ssDNA and GA-ssDNA are same as in figure 2 of main text.

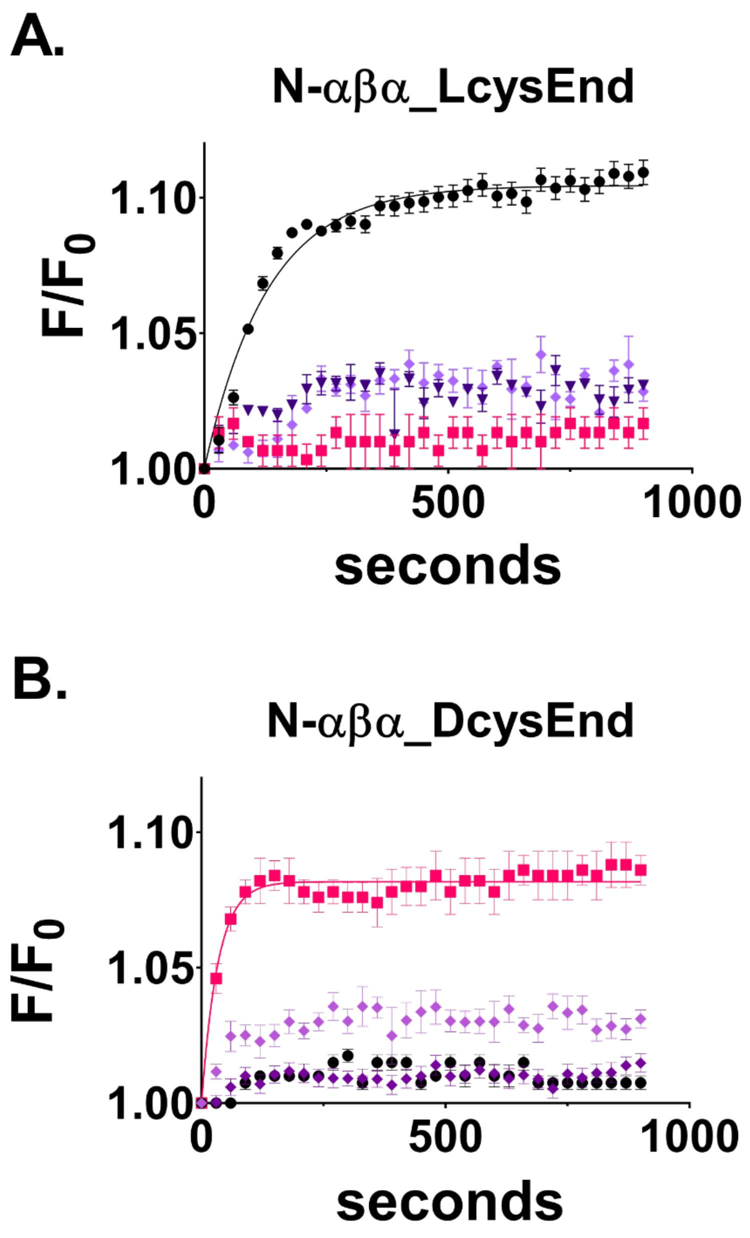


**Figure S8: Comparative adsorption profiles of synthetic N-αβα prototypes to TC-ssDNA and fluorescein dye. A.** Adsorption isotherms of N-αβα_LcysEnd and **B.** N-αβα_DcysEnd binding to TC-ssDNA and fluorescein dye for UP and DOWN orientations of the magnet. *TC-ssDNA + UP (black)*, *TC-ssDNA + DOWN (pink)*, *fluorescein + UP (purple)*, *fluorescein + DOWN (violet)*. Data were plotted as $F/F_0$ vs seconds and fitted to standard one-phase association equation using GraphPad Prism software. Error bars represent standard deviation from three to four independent measurements. Adsorption curves for N-αβα_LcysEnd and N-αβα_DcysEnd are same as in figure 3 of main text.

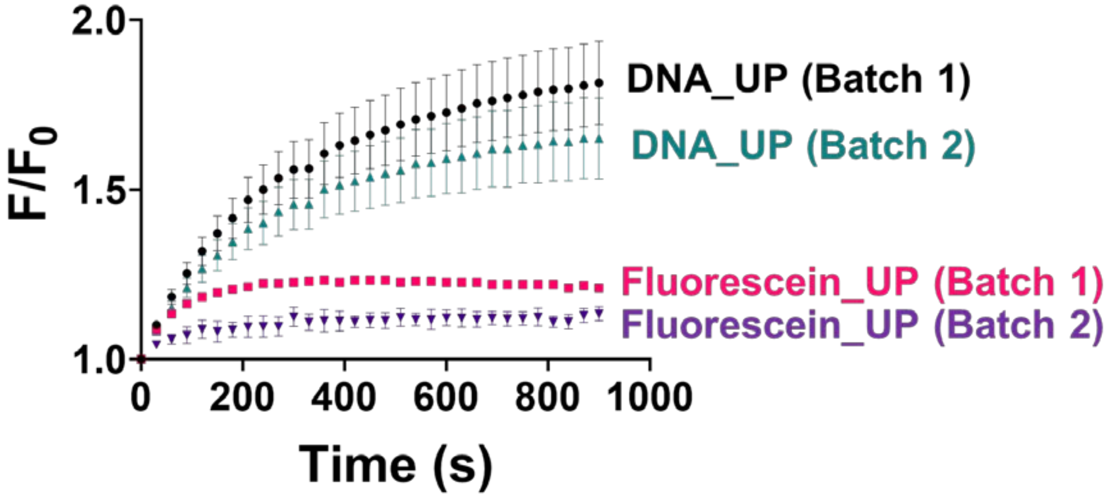


**Figure S9:** Binding of TC-ssDNA to N-αβα prototype for two independent batches of protein purification. Shown here are binding curves of TC-ssDNA and fluorescein (i.e., digested DNA) to adsorbed N-αβα prototype for UP orientation, as in Figure 2 of main text, for two independent batches of purification rounds. Batch 1 traces are same as in Figure 2A of main text.

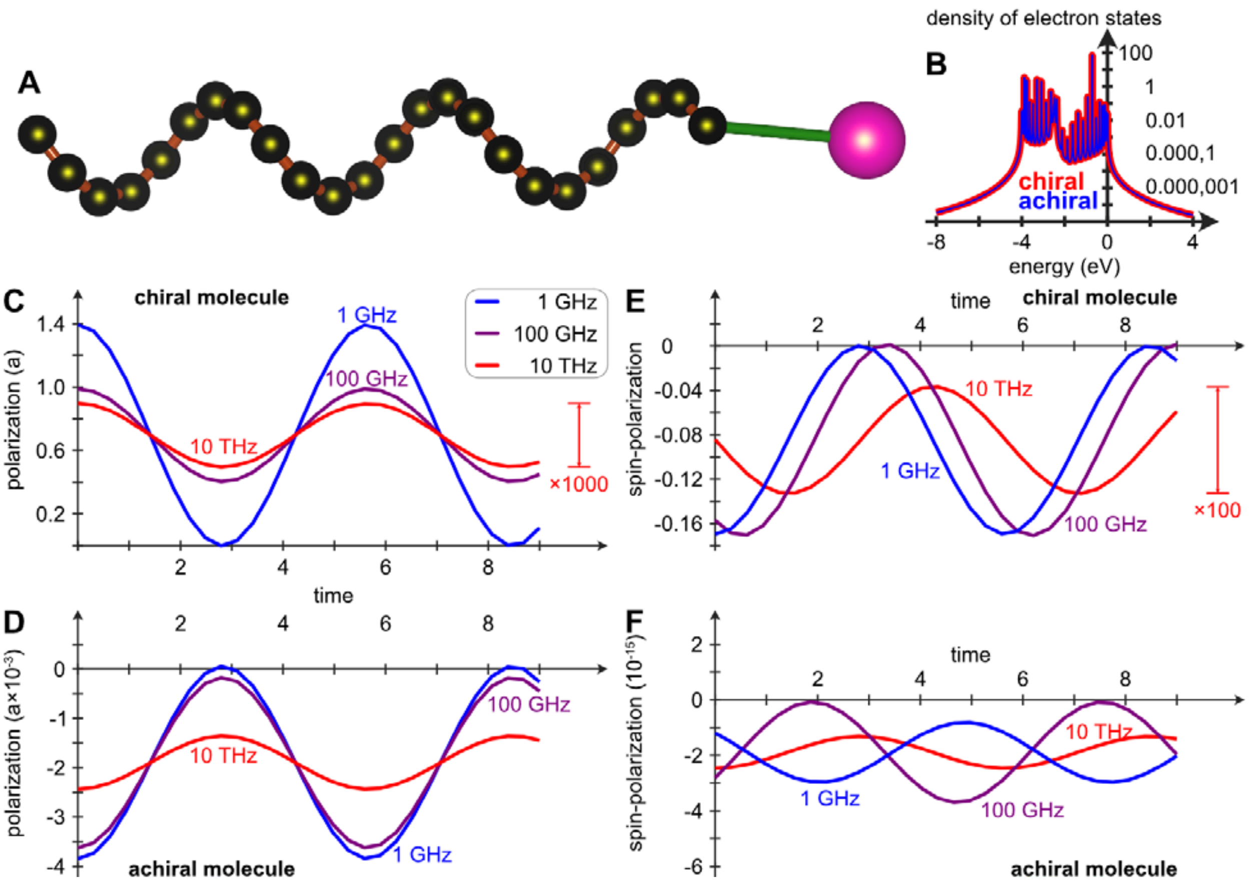


**Figure S10: A.** Schematic of the chiral system (black balls, brown bonds) connected to the external charge (magenta ball, green coupling). The corresponding achiral system is represented by a zig-zag chain of sites. **B**. Density of electron states for the chiral (red) and achiral (blue) molecules. The chemical potential is set to 0. **C**. & **D**. Time-dependent variations of the dominant component of the electric polarization vector $\boldsymbol{P}$ for the **C**. chiral and **D**. achiral system, under the time-dependent coupling between the system and external charge. The time-dependent coupling varies harmonically with frequencies $\nu_0 = 1$ GHz, 100 GHz, 10 THz. Results with more frequencies are presented in the supplementary information. **E**. & **F**. Time-dependent variations of the spin-polarizations corresponding to the polarizations in **C**. & **D**. The sites in the system host electron levels at $\epsilon_m = -2t_0$ and are elastically coupled by the nearest neighbor coupling $t_0 = 1$ eV and next-nearest neighbor coupling $\lambda_0 = 1$ meV, whereas the inelastic coupling strengths are $t_1 = t_0/10$ and $\lambda_1 = \lambda_0/10$. The system vibrates with the frequency corresponding to the energy $\omega_0 = 0.5$ eV, the helix radius a $= 5$ Å, and the temperature is set to $T = 300$ K.

**Table S1: Coding DNA sequences of P-loop prototypes used in this study.**

| **Protein** | **Coding DNA sequence** |
|---|---|
| N-αβα (ref. [4]) | ATGACTCGTGATGATGCAAAACGTGTAGCGGAAGAAGCGA AGCGTCGCGGTGTTGGTAGCGGCCGTGTGATTATCGTAATT GTGGGTCCAAGCGGCGCAGGCAAAACCACCCTGCTCGAAC TGGCTAAAGAAGCTAAGAAGGAGGTGTGGCTCGAGCACCA CCACCACCATCAC**TGA** |
| N-αβα with terminal cysteine (this study) | ATGACTCGTGATGATGCAAAACGTGTAGCGGAAGAAGCGA AGCGTCGCGGTGTTGGTAGCGGCCGTGTGATTATCGTAATT GTGGGTCCAAGCGGCGCAGGCAAAACCACCCTGCTCGAAC TGGCTAAAGAAGCTAAGAAGGAGGTGTGGCTCGAGCACCA CCACCACCATCAC**TGCTGA** |
| N-βα (ref. [4]) | ATGCGTGTGATTATCGTAATTGTGGGTCCAAGCGGCGCAGG CAAAACCACCCTGCTCGAACTGGCTAAAGAAGCTAAGAAG GAGGTGTGGCTCGAGCACCACCACCACCATCAC**TGA** |
| N-βα with terminal cysteine (this study) | ATGCGTGTGATTATCGTAATTGTGGGTCCAAGCGGCGCAGG CAAAACCACCCTGCTCGAACTGGCTAAAGAAGCTAAGAAG GAGGTGTGGCTCGAGCACCACCACCACCATCAC**TGCTGA** |

The DNA sequences of N-αβα and N-βα prototype are shown from reference [4]. The DNA sequence codon for the incorporated cysteine is shown in bold red. The terminal stop codon is shown in bold.

**Table S2: Amino acid sequences of protein constructs used in this study.**

| Prototype | Amino acid sequence | Molecular weight (kDa) |
|---|---|---|
| **Prototypes expressed in purified from BL.21 E. coli cells** | | |
| N-αβα (ref. [4]) | MTRDDAKRVAEEAKRRGVGSGRVIIVIV**GPSG AGKT**TLLELAKEAKKEV*WLEHHHHHH* | 6.4 |
| N-αβα with terminal cysteine | MTRDDAKRVAEEAKRRGVGSGRVIIVIV**GPSG AGKT**TLLELAKEAKKEV*WLEHHHHHH***C** | 6.5 |
| N-βα (ref. [4]) | MRVIIVIV**GPSGAGKT**TLLELAKEAKKEV*WLE HHHHHH* | 4.3 |
| N-βα with terminal cysteine | MRVIIVIV**GPSGAGKT**TLLELAKEAKKEV*WLE HHHHHH***C** | 4.4 |
| **Chemically synthesized constructs** | | |
| N-αβα_LcysEnd | MTRDDAKRVAEEAKRRGVGSGRVIIVIV**GPSG AGKT**TLLELAKEAKKEV**C** | 5.3 |
| N-αβα_DcysEnd | MTRDDAKRVAEEAKRRGVGSGRVIIVIV**GPSG AGKT**TLLELAKEAKKEV**C** | 5.3 |
| *L*-Precursor | CSIERIRRASVEELTEV**PGIGP**RLARRILERL | 3.7 |
| *D*-Precursor | CSIERIRRASVEELTEV**PGIGP**RLARRILERL | 3.7 |

Bacterially expressed and purified prototypes contained a C-terminal expression tag that included a tryptophan (W) residue to allow determination of protein concentration by absorbance at 280 nm, and a 6xHis tag for purification (annotated in italics). Terminal cysteine residues are shown in bold red. For synthetic N-αβα_DcysEnd, the terminal cysteine which is in the D-form is shown in underlined red. P-loop motif is shown in bold in all prototypes. For synthetic HhH peptides, the conserved PGIGP motif is shown in bold. The HhH peptide with *D*-form amino acids is shown in red. Molecular weight for each construct was calculated from the amino-acid sequence using ExPASy ProtParam tool [5]

**Table S3: DNA oligonucleotides used in this study**

| Oligos | Sequence (5' to 3') | Figure panels from main text |
|---|---|---|
| TC-ssDNA | 6-FAM - TACTTCTCTTCTCTCTCCTCGACT | 2A, 2C, 3A, 3B, 4B |
| GA-ssDNA | 6-FAM - AGTCGAGGAGAGAGAAGAGAAGTA | 2B, 2C |
| $DNA_{12}$-sense | 6-FAM - TAGATCGATCGC | 3D, 3E |
| $DNA_{12}$-antisense | GCGATCGATCTA | 3D, 3E |

6-FAM = 6-Carboxyfluorescein

**Table S4: Binding properties of N-αβα prototype to ssDNA constructs as measured by fluorescence anisotropy.**

| ssDNA constructs | $K_D$ (μM) | h | $R^2$ |
|---|---|---|---|
| TC-ssDNA | 0.67 (±0.16) | 3.92 (±0.64) | 0.95 |
| GA-ssDNA | 1.70 (±0.68) | 2.98 (±2.44) | 0.83 |

Data is reproduced from ref. [4]. Values in parenthesis represents standard deviation from four to eight independent experiments.
